# Multi-omics analysis of sarcospan overexpression in *mdx* skeletal muscle reveals compensatory remodeling of cytoskeleton-matrix interactions that promote mechanotransduction pathways

**DOI:** 10.1101/2022.07.26.501621

**Authors:** Jackie L. McCourt, Kristen M. Stearns-Reider, Hafsa Mamsa, Cynthia Shu, Mohammad Hossein Afsharinia, Elizabeth M. Gibbs, Kara M. Shin, Yerbol Z. Kurmangaliyev, Lauren R. Schmitt, Kirk C. Hansen, Rachelle H. Crosbie

## Abstract

**Background:** The dystrophin-glycoprotein complex (DGC) is a critical adhesion complex of the muscle cell membrane, providing a mechanical link between the extracellular matrix (ECM) and the cortical cytoskeleton that stabilizes the sarcolemma during repeated muscle contractions. One integral component of the DGC is the transmembrane protein, sarcospan (SSPN). Overexpression of SSPN in the skeletal muscle of *mdx* mice (murine model of DMD) restores muscle fiber attachment to the ECM in part through an associated increase in utrophin and integrin adhesion complexes at the cell membrane, protecting the muscle from contraction-induced injury. In this study, we utilized transcriptomic and ECM protein-optimized proteomics data sets from wild-type, *mdx*, and *mdx* transgenic (*mdx* ^TG^) skeletal muscle tissues to identify pathways and proteins driving the compensatory action of SSPN overexpression.

**Methods:** The tibialis anterior and quadriceps muscles were isolated from wild-type, *mdx*, and *mdx* ^TG^ mice and subjected to bulk RNA-Seq and global proteomics analysis using methods to enhance capture of ECM proteins. Data sets were further analyzed through the Ingenuity Pathway Analysis (QIAGEN) and integrative gene set enrichment to identify candidate networks, signaling pathways, and upstream regulators.

**Results:** Through our multi-omics approach, we identified 3 classes of differentially expressed genes and proteins in *mdx* ^TG^ muscle, included those that were: 1) unrestored (significantly different from wild-type, but not from *mdx*), 2) restored (significantly different from *mdx*, but not from wild-type), and 3) compensatory (significantly different from both wild-type and *mdx*). We identified signaling pathways that may contribute to the rescue phenotype, most notably cytoskeleton and ECM organization pathways. ECM optimized-proteomics revealed an increased abundance of collagens II, V, and XI, along with β-spectrin in *mdx* ^TG^ samples. Using Ingenuity Pathway Analysis, we identified upstream regulators that are computationally predicted to drive compensatory changes, revealing a possible mechanism of SSPN rescue through a rewiring of cell-ECM bidirectional communication. We found that SSPN overexpression results in upregulation of key signaling molecules associated with regulation of cytoskeleton organization and mechanotransduction, including Rho, RAC, and Wnt.

**Conclusions:** Our findings indicate that SSPN overexpression rescues dystrophin deficiency partially through mechanotransduction signaling cascades mediated through components of the ECM and the cortical cytoskeleton.

## BACKGROUND

Duchenne muscular dystrophy (DMD) is a progressive muscle wasting disorder caused by mutations in the *DMD* gene encoding the protein dystrophin (1). While dystrophin is expressed in multiple tissues, loss of dystrophin in the context of DMD is particularly detrimental to skeletal and cardiac muscle function (2). Dystrophin is a component of the dystrophin-glycoprotein complex (DGC) that stabilizes the muscle fiber sarcolemma and mediates the linkage between the extracellular matrix (ECM) and the intracellular actin cytoskeleton (3–6). Loss of dystrophin in DMD results in the absence of the DGC, destabilizing the sarcolemma and rendering it susceptible to contraction-induced damage (7). Over time, chronic injury caused by asynchronous cycles of myofiber degeneration and regeneration results in failed muscle regeneration, inflammation, and replacement of functional muscle fibers with fibrosis (8). Patients with DMD present with high plasma levels of muscle creatine kinase at birth, muscle fibrosis and hypertrophy, weakness of the proximal muscles, loss of ambulation, and pulmonary and cardiac dysfunction leading to premature death in the 2^nd^ to 3^rd^ decade of life (9–11). While there are some FDA approved treatment options for DMD, including corticosteroids and exon skipping drugs that slow progression of the disease, there is still no cure.

The dystrophin-deficient *mdx* mouse is the most widely used mouse model for DMD as it exhibits much of the pathology observed in patient skeletal muscle, albeit much milder, including elevated creatine kinase levels, increased levels of degeneration and regeneration, fibrosis, and reduced grip strength and whole-body tension (12,13). While many therapeutic genes that improve sarcolemma instability, decrease fibrosis, and increase force in *mdx* muscle have been identified (14–21), there are gaps in understanding whether their protective mechanisms affect overlapping molecular processes and signaling pathways. Furthermore, there is emerging evidence of an expanded network of DGC interacting proteins, suggesting that it functions as a central unit to integrate cellular signaling, lateral force transmission, ion channel function, and cytoskeletal organization (22,23).

Sarcospan (SSPN), a core component of the DGC (24–26) prevents muscle degeneration and histopathology in *mdx* mice in a mechanism that is dependent on increased abundance of utrophin and integrin α7β1 at the sarcolemma (27–31). Together with the sarcoglycans (SG), SSPN forms a tight subcomplex (26) that anchors α-dystroglycan, a receptor for many ECM proteins (32), to the cell membrane to stabilize the cytoskeleton-matrix linkage (33). SSPN increases SG protein levels and restores integrity of the SG-SSPN subcomplex at the sarcolemma in *mdx* skeletal and cardiac muscles (27,28,30,31,34,35). SSPN overexpression in *mdx* mice also increased abundance of dystroglycan, enhanced matriglycan glycosylation of α-dystroglycan, and improved laminin binding (27,30,34,36). Using genetic approaches, we previously demonstrated that SSPN function is dependent on matriglycan modification of α-dystroglycan, which directly interacts with laminin (34). Restoration of cell-matrix interactions by SSPN improved post-exercise activity, protected against eccentric contraction-induced force loss, and prevented declines in pulmonary and cardiac functions (30,31,35).

In the current study, we interrogated the effects of SSPN overexpression on the *mdx* transcriptome and proteome. Our findings build on many prior studies including microarray data sets (Omnibus GSE465, GSE1004, and GSE1007) (37) and proteomic analysis (37–44) of *mdx* muscle and DMD patient muscle biopsies, which revealed signaling pathways that contribute to the dystrophic pathology including anaerobic metabolism, cytoskeleton remodeling, calcium handling, adipogenesis, fibrosis, and endoplasmic reticulum stress. Given that there are many emerging approaches to treating DMD (dystrophin-dependent as well as dystrophin-independent), determining the effects of such treatments on cellular and molecular processes is important for therapeutic design. Such studies also have the potential to reveal overlapping pathways that are critical for prevention of muscle degeneration as well as pathways that are unaltered by a particular therapy (thereby informing molecular processes that are not major contributors to pathology). For instance, therapeutic restoration of dystrophin expression in *mdx* muscle by anti-sense oligonucleotide exon skipping normalizes protein and mRNA expression toward wild-type levels, but does not affect the expression of microRNAs (37). In our study, we found that SSPN overexpression in *mdx* muscle does not restore all transcripts and proteins to wild-type levels but ameliorates muscle pathology through compensatory changes in ECM and cytoskeletal composition, including key signaling molecules associated with regulation of cytoskeleton organization and mechanotransduction.

## METHODS

### Mouse models

Wild-type (C57BL/6J) and *mdx* mice were purchased from Jackson Laboratories (Bar Harbor, ME, USA). We have generated and extensively characterized multiple lines of SSPN-transgenic *mdx* mice expressing either human SSPN (hSSPN, line 3) and murine SSPN (mSSPN, line 28) under the control of the human skeletal α-actin promoter (27,30,36). hSSPN and mSSPN exhibit a high degree of identity (>85%) at the amino acid level, effectively ameliorate *mdx* pathology, and were included in the study with the rationale that overlapping findings from different transgenic lines would strengthen their relevance. All mouse colonies were granted by the UCLA Animal Welfare Assurance (approval #A3196-01).

### Poly-A enriched RNA sequencing

Liquid nitrogen frozen tibialis anterior muscles from 12-week-old male wild-type, *mdx*, and *mdx* ^TG^ (line 28) were pulverized in a mortar and pestle cooled by liquid nitrogen, and homogenized into TRIzol (Thermo Fisher Scientific, Waltham, MA) using syringe homogenization with a 21g needle. RNA was extracted using TRIzol phase separation followed by QIAGEN RNeasy column purification using the manufacturer’s instructions (QIAGEN, Hilden, Germany), followed by DNase treatment. RNA concentration and quality were measured using the TapeStation 4200 (Agilent Technologies, Santa Clara, CA). RNA-Seq libraries were prepared from total RNA using the TruSeq Stranded mRNA Library Prep Kit (Illumina, San Diego, CA). High-throughput sequencing was performed at UCLA Technology Center for Genomics & Bioinformatics using the Illumina HiSeq 4000 platform (paired-end 75 bp reads). Demultiplexing was performed with Illumina Bcl2fastq2 v 2.17 program. The total numbers of sequenced reads were 53-67 million per sample. RNA-Seq reads were mapped to the mouse reference genome (mm10) using STAR (45). For each sample, 86% of reads were uniquely mapped to the genome. Expression levels were quantified for annotated genes (Ensembl v.92) and raw gene counts were normalized to CPM values (counts-per-million). The analysis was focused on genes with CPM > 1 in at least two samples (13846 genes). Differential expression analysis was performed using edgeR-QLF (46). Differentially expressed genes (DEGs) were identified at 1% FDR and fold-change > 2.

### Mass Spectrometry

The mass spectrometry dataset was first published in part in Stearns-Reider *et al* (47). Quadriceps muscles were harvested from 20-week-old male wild-type, *mdx*, and *mdx* ^TG^ (line 3) mice and snap frozen in liquid nitrogen. Samples were then prepared for mass spectrometry analysis as previously described (48). In short, samples were pulverized in liquid nitrogen and lyophilized. For each sample, 5 mg (dry weight) of tissue was homogenized in 200mL/mg high-salt buffer (HS buffer) containing a 1x protease inhibitor (48). Following three rounds of HS buffer wash, pellets were treated with 6M guanidine extraction buffer. The remaining pellets from each tissue, representing insoluble ECM proteins, were digested with freshly prepared hydroxylamine buffer, as previously described (49). 100μl of the cellular fraction (combined fractions 1, 2, and 3) and 200μl of the soluble and insoluble ECM fractions were enzymatically digested with trypsin using a filter-aided sample prep approach and C18 tip cleanup. Samples were then analyzed by liquid chromatography – data dependent acquisition tandem mass spectrometry (LC-MS/MS), as previously described (50). Samples were analyzed on a Q Exactive HF Orbitrap mass spectrometer (Thermo Fisher Scientific) coupled to an EASY-nanoLC 1000 system through a nanoelectrospray source. The analytical column (100 μm i.d. × 150 mm fused silica capillary packed in house with 4 μm 80 Å Synergi Hydro C18 resin (Phenomenex; Torrance, CA)) was then switched online at 600 nL/min for 10 minutes to load the sample. The flow rate was adjusted to 400 nL/min, and peptides were separated over a 120-min linear gradient of 2–40% ACN with 0.1% FA. Data acquisition was performed using the instrument supplied Xcalibur (Thermo Fisher Scientific, San Jose, CA) software in positive ion mode. MS/MS spectra were extracted from raw data files and converted into mgf files using a PAVA script (University of California, San Francisco, MSF, San Francisco, CA). These mgf files were then independently searched against mouse SwissProt database using an in-house Mascot server (version 2.2.06; Matrix Science, London, UK). Mass tolerances were +/-10 ppm for MS peaks and +/-0.5 D for MS/MS fragment ions. Trypsin specificity was used allowing for one missed cleavage. Met oxidation, Pro oxidation, protein N-terminal acetylation, and peptide N-terminal pyroglutamic acid formation were allowed for variable modifications, whereas carbamidomethyl of Cys was set as a fixed modification. Following Mascot searches, data was directly loaded into ScaffoldTM (Proteome Software Inc.). Peptide spectral matches were directly exported with a 99% confidence in protein identifications and at least 2 unique peptides per protein, resulting in a false discovery rate of 0.54%. Two-group comparisons were done by two-tailed Student’s t-tests. Partial least squares-discriminant analysis (PLSDA) was performed using MetaboAnalyst (version 3.0) with sum and range scaling normalizations.

### Indirect Immunofluorescence Staining of Muscle Sections

Transverse cryosections (10μm) from the quadriceps muscles of wild-type, *mdx*, and *mdx* ^TG^ (line 3) mice at 5 months of age were incubated in blocking buffer (3% BSA in phosphate buffered saline (PBS)) for 30 minutes at room temperature. Avidin/biotin blocking kit (Vector Laboratories, Newark, CA) was used according to manufacturer’s instructions. Sections were then incubated with the following primary antibodies diluted 1:200 in PBS overnight at 4°C: β-spectrin non-erythrocyte (hpa012685, Sigma, St. Louis, MO), collagen type II (ab34712, Abcam, Waltham, MA), collagen type V (ab7046, Abcam, Waltham, MA). Sections were washed in PBS for 3 x 30 min at room temperature. Primary antibodies were detected by biotinylated anti-rabbit (BA-1000; 1:500; Vector Laboratories, Newark, CA). Fluorescein-conjugated avidin D (A-2001; 1:500; Vector Laboratories) was used to detect secondary antibodies. Sections were mounted in Vectashield (Vector Laboratories, Newark, CA) and imaging was performed using a Zeiss Axio Imager M2 (Carl Zeiss Inc., Thornwood, NY) with a Hammatsu ORCA-Flash 4.0 V3 digital complementary metal oxide semiconductor camera and a Plan-Aprochromat 20×/0.8 M27 objective.

### Gene Ontology and Ingenuity Pathway Analysis

Gene ontology (GO) enrichment analysis for RNA sequencing and proteomic data sets was performed using the PANTHER Classification System online user interface (51) and the statistical overrepresentation test. Fold enrichment values and p-values from the over-representation test were plotted using GraphPad Prism. RNA sequencing and proteomic data sets were also analyzed through the use of Ingenuity Pathway Analysis (QIAGEN) for generating networks and upstream regulators(52). Upstream regulators were sorted by activation score and the top five inhibited and activated regulators were reported with corresponding target genes identified in the data sets.

### Integrated Gene Set Enrichment Analysis

Significantly differentially expressed transcripts and proteins between *mdx* and *mdx* ^TG^ were pooled as input for integrated gene set enrichment analysis using Database for Annotation, Visualization and Integrated Discovery (DAVID, version 2021). Each molecule was categorized as unrestored (significantly different from wild-type, not significantly different from *mdx*), restored (significantly different from *mdx*, not significantly different from wild-type), or compensatory (significantly different from both wild-type and *mdx*). Data were visualized on Cytoscape version 3.9.1.

### Supplemental Figure Methods

For a systems level network analysis, DEGs from the three identified categories (restored, unrestored, compensatory) were analyzed using the Database for Annotation, Visualization and Integrated Discovery platform (DAVID, version 2021). This analysis was also performed with the proteomic data set. Gene Ontology (GO) terms with corresponding parameter thresholds of p < 0.05 and minimum gene count = 2 were used as input and visualization in Cytoscape with the EnrichmentMap plugin.

## RESULTS

### Gene Expression and Proteomic Analyses Reveal Compensatory and Restored Pathways

In the current study, our goal was to interrogate the effects of SSPN overexpression on gene and protein expression in *mdx* muscle relative to wild-type and *mdx* (non-transgenic) controls. Gene expression analysis was performed using traditional poly-A enriched RNA sequencing of tibialis anterior muscles isolated from 12-week-old wild-type, *mdx*, and *mdx* ^TG^ mice. Principal component analysis of the sequencing data reveals clear clustering of each genotype (Fig. 1a). In comparing paired genotypes, we identified 1073 DEGs (853 upregulated, 220 downregulated) in wild-type versus *mdx* muscle and 748 DEGs (471 upregulated, 277 downregulated) in *mdx* relative to *mdx* ^TG^ samples. The largest difference in the transcriptomic profile was evident between wild-type and *mdx* ^TG^ with 1857 DEGs (1306 upregulated, 551 downregulated). Heatmap analysis of DEGs in *mdx* ^TG^ relative to controls (Fig. 1b) reveals patterns that can be categorized as: 1) unrestored (significantly different from wild-type, not significantly different from *mdx*), 2) restored (significantly different from *mdx*, not significantly different from wild-type), and 3) compensatory (significantly different from both wild-type and *mdx*).

**Figure 1.**
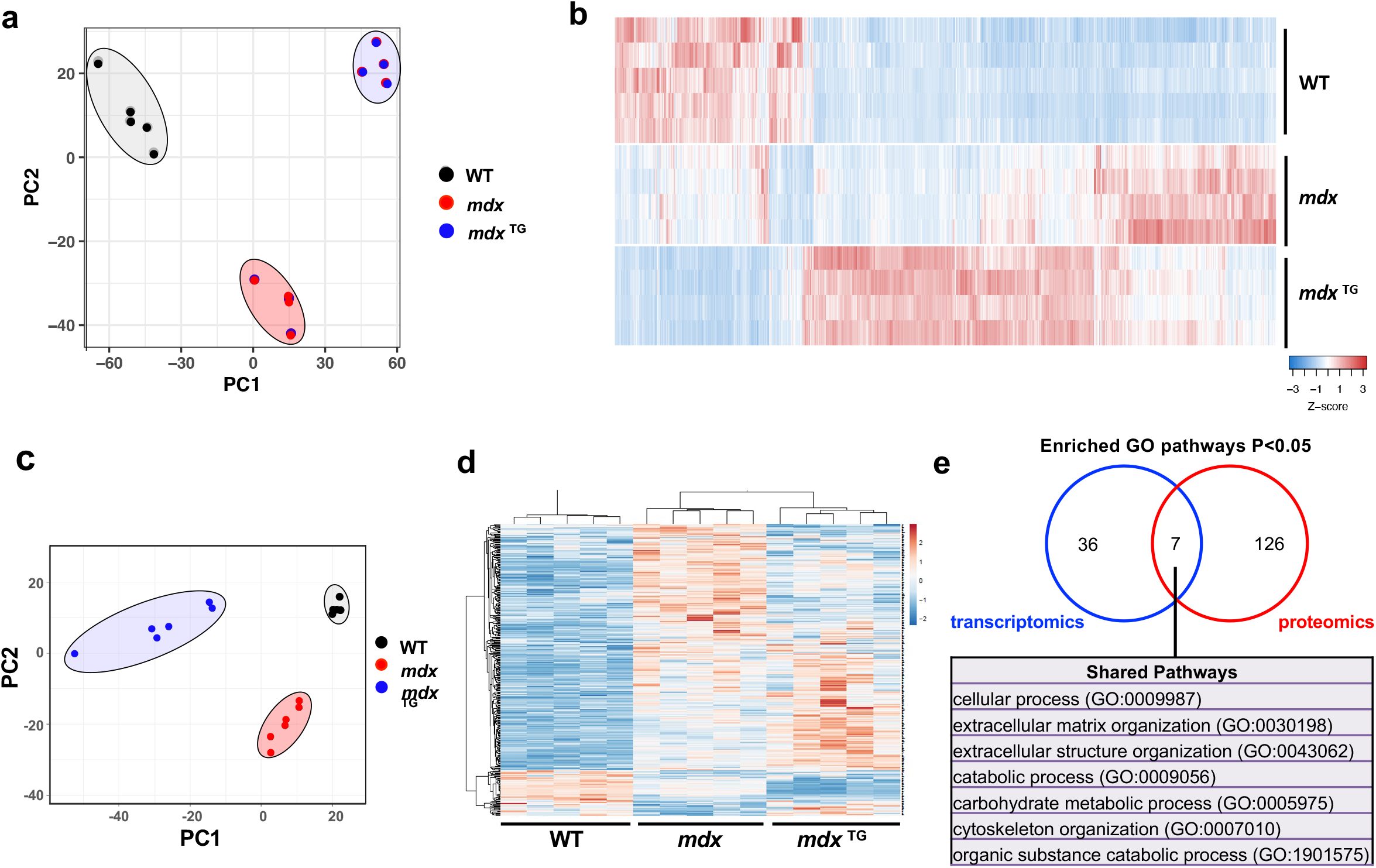
Overview of RNA sequencing and mass spectrometry reveals distinct transcriptomic and proteomic profiles of SSPN overexpression rescue. **a**) Principal component analysis (PCA) of RNA sequencing data in wild-type (WT), *mdx*, and mSSPN transgenic (*mdx* ^TG^) tibialis anterior muscle at 12 weeks of age. **b)** Heat map of DEGs. **c)** PCA of mass spectrometry data from WT, *mdx*, and hSSPN transgenic (*mdx* ^TG^) quadriceps muscle at 20 weeks of age. **d**) Heat map of differentially expressed proteins (DEPs). **e**) Overlap of Gene Ontology (GO) terms enriched in WT vs *mdx* ^TG^ in transcriptomics vs proteomics data using PANTHER GO analysis platform.

Given the effects of SSPN overexpression on DEGs, we next investigated the influence of SSPN on protein expression utilizing a mass spectrometry approach that improves capture of typically insoluble ECM proteins (47,53). From our proteomics analysis of wild-type, *mdx*, and *mdx* ^TG^ quadriceps muscle, we identified a total of 1679 proteins in all three genotypes. Principal component analysis revealed distinct clustering of the wild-type, *mdx*, and *mdx* ^TG^ samples (Fig. 1c). Similar to the RNA sequencing data, we found sets of proteins in restored, unrestored, and compensatory categories (Fig. 1d).

### Pathway Enrichment Analysis Identifies Compensatory Changes in Fibrillar Collagens and Components of the Actin Cytoskeleton

Comparative analysis of enriched GO pathways in the transcriptomic and proteomics data sets revealed seven shared pathways, including those associated with ECM organization, extracellular structure organization, and cytoskeleton organization (Fig. 1e). By separating enriched GO pathways into the compensatory, restored, and unrestored subcategories for both transcriptomic and proteomic data sets, we observed additional pathways of interest such as those associated with calcium ion binding, NAD/NADP binding, ubiquitin ligase binding, and oxidoreductase binding (Supplemental Figure 1). Given the identification of the broader categories of ECM organization and cytoskeleton organization identified in the *mdx* ^TG^ model through GO pathway analysis, we curated lists of actin cytoskeleton- and ECM-associated genes in the RNA sequencing data set based on GO categories and subsequently visualized this data using traditional heat maps. The actin cytoskeleton and ECM genes clustered into restored, unrestored, and compensatory categories. It is noteworthy that the compensatory genes represent the largest category of DEGs in this comparison (Fig. 2a-b).

**Figure 2.**
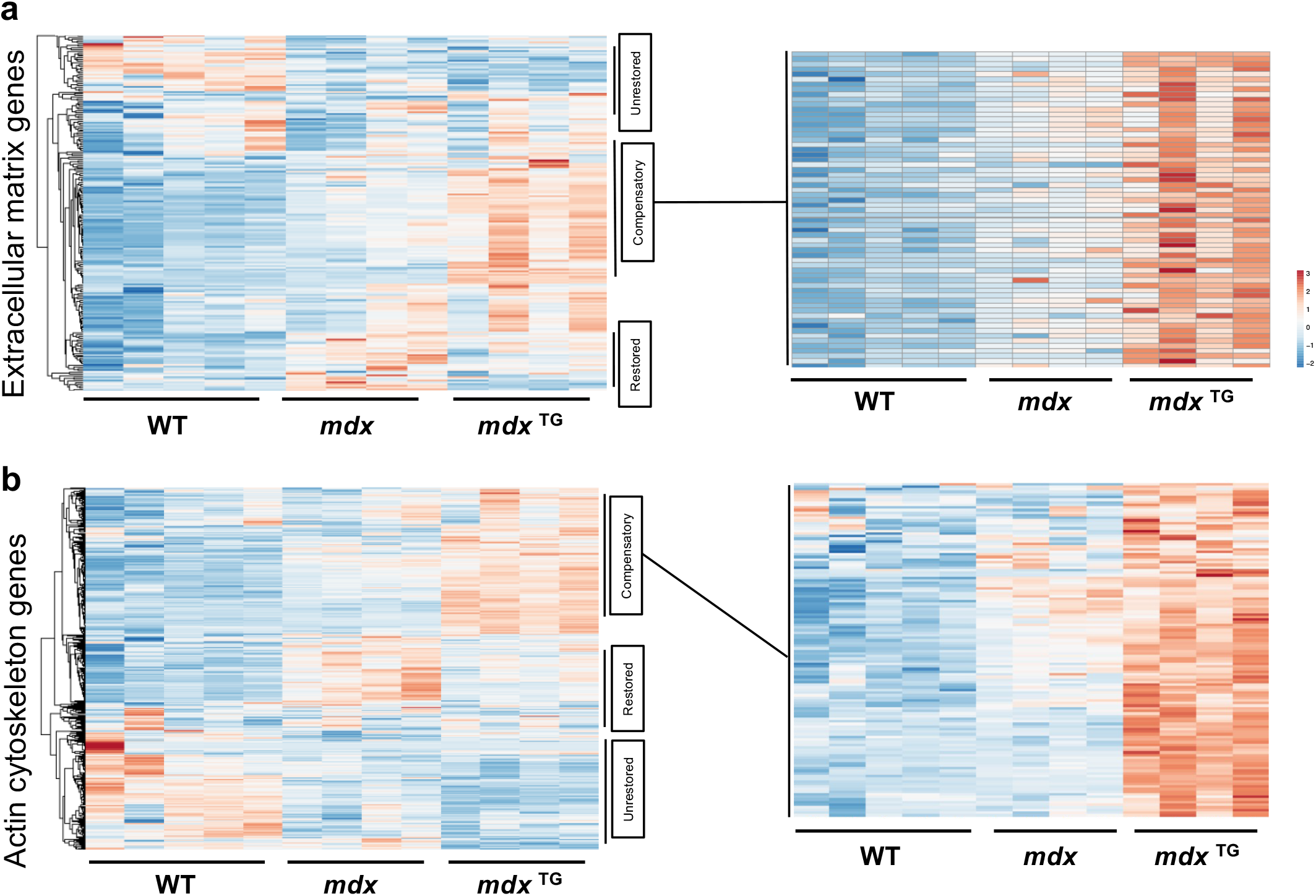
SSPN overexpression results in compensatory upregulation of ECM and actin cytoskeleton genes. Heat maps of GO-term curated ECM genes (**a**) and actin cytoskeleton genes (**b)** from RNA sequencing data each with restored, unrestored, and compensatory expression patterns in the *mdx* ^TG^ muscle. Expanded heat map insets emphasize that compensatory expression patterns are overwhelmingly from upregulated genes in the *mdx* ^TG^ muscle.

We next analyzed the expression of ECM and cytoskeletal proteins from the mass spectrometry data set. ECM proteins were classified into 8 functional categories, including: 1) structural ECM, 2) other ECM, 3) network collagen, 4) matricellular, 5) fibrillar collagen, 6) fibril-associated collagens with interrupted triple helices (FACIT), 7) ECM regulator, and 8) basement membrane proteins (Fig. 3a). Cytoskeletal proteins were classified into 9 functional categories, including: 1) actins and microfilaments, 2) actin-associated, 3) tubulins, 4) microtubule associated, 5) annexins, 6) intermediate filaments, 7) spectrins, 8) myosins, and 9) sarcomere-associated proteins (Fig. 3b). ECM regulators and fibrillar collagens were upregulated in *mdx* ^TG^ compared to both wild-type and *mdx* muscles (Fig. 3a). For cytoskeletal proteins, we observed significant downregulation in actins, microfilaments, and actin-associated proteins, as well as decreased abundance of myosins and other sarcomere-associated proteins such as tropomyosins and troponin I and T (Fig. 3b). Abundance of microtubule associated proteins and spectrins was increased in *mdx* ^TG^ samples relative to wild-type controls. Within each functional classification, we additionally identified several proteins with increased expression in *mdx* ^TG^ muscle compared to both wild-type and *mdx* muscle including cathepsins and integrins (Fig. 4a), fibrillar collagens II, V, and XI (Fig. 4b), and non-erythrocyte β-spectrin (Sptbn1, Spectrins, Fig. 4c). Interestingly, some actin isoforms and actin-associated proteins were decreased in *mdx* ^TG^ muscle compared to both wild-type and *mdx* muscle including skeletal and cardiac α-actins (Acta1, Actc1) and α-actinin (Actn3, Fig. 4d).

**Figure 3.**
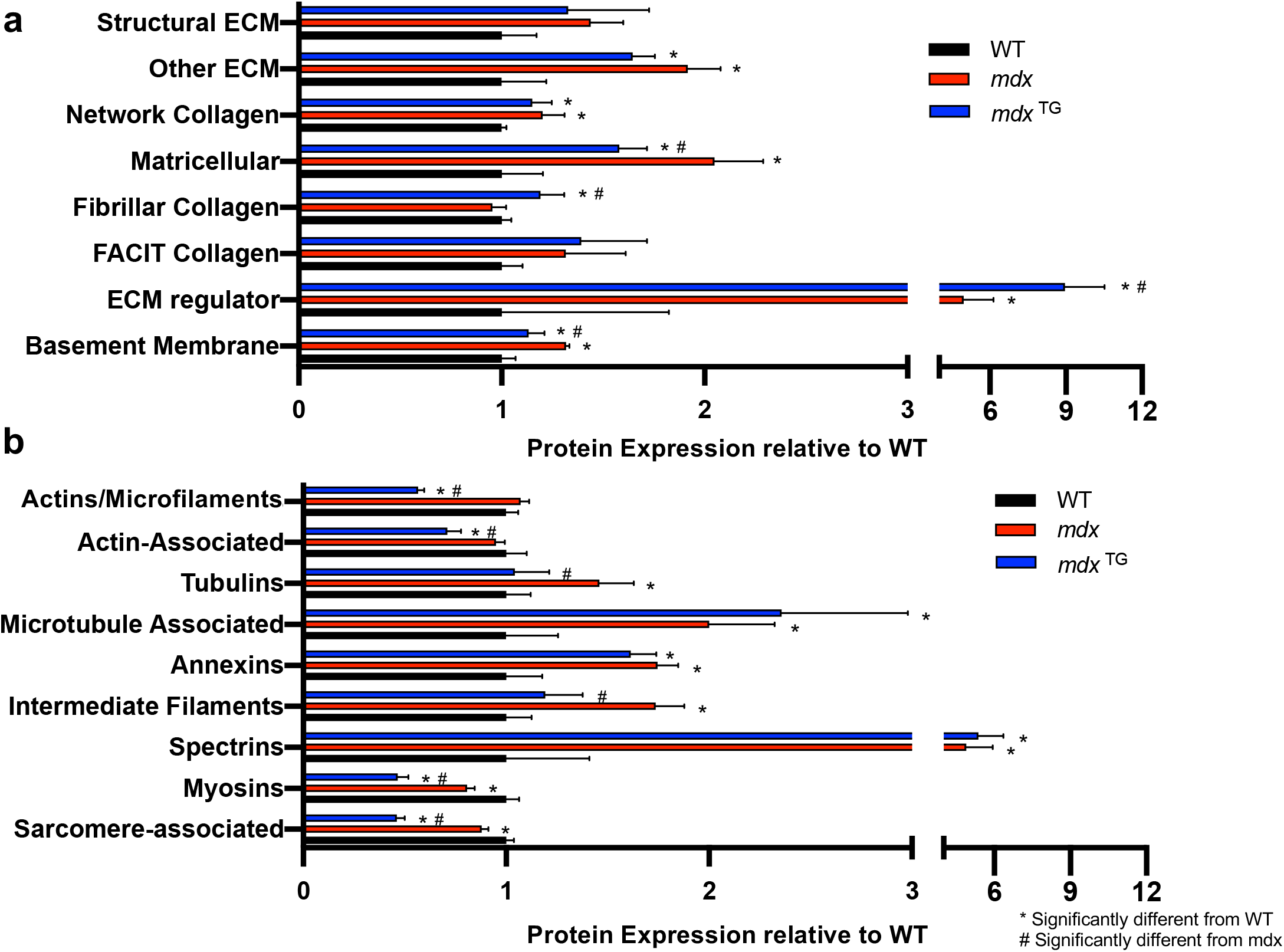
Compensatory changes in functional classes of ECM and cytoskeletal proteins in *mdx* ^TG^ muscle. **a**) Graph of the abundance of ECM proteins in 8 primary categories relative to WT. **b**) Graph of the abundance of cytoskeletal proteins in 9 primary categories relative to WT (*p<0.05 compared to WT, # p<0.05 compared to *mdx* by unpaired t-test). By category, *mdx* ^TG^ muscle had compensatory expression (both significantly different from WT and *mdx*) of matricellular, ECM regulator, basement membrane, actins/microfilaments, actin-associated, myosins, and sarcomere-associated proteins.

**Figure 4.**
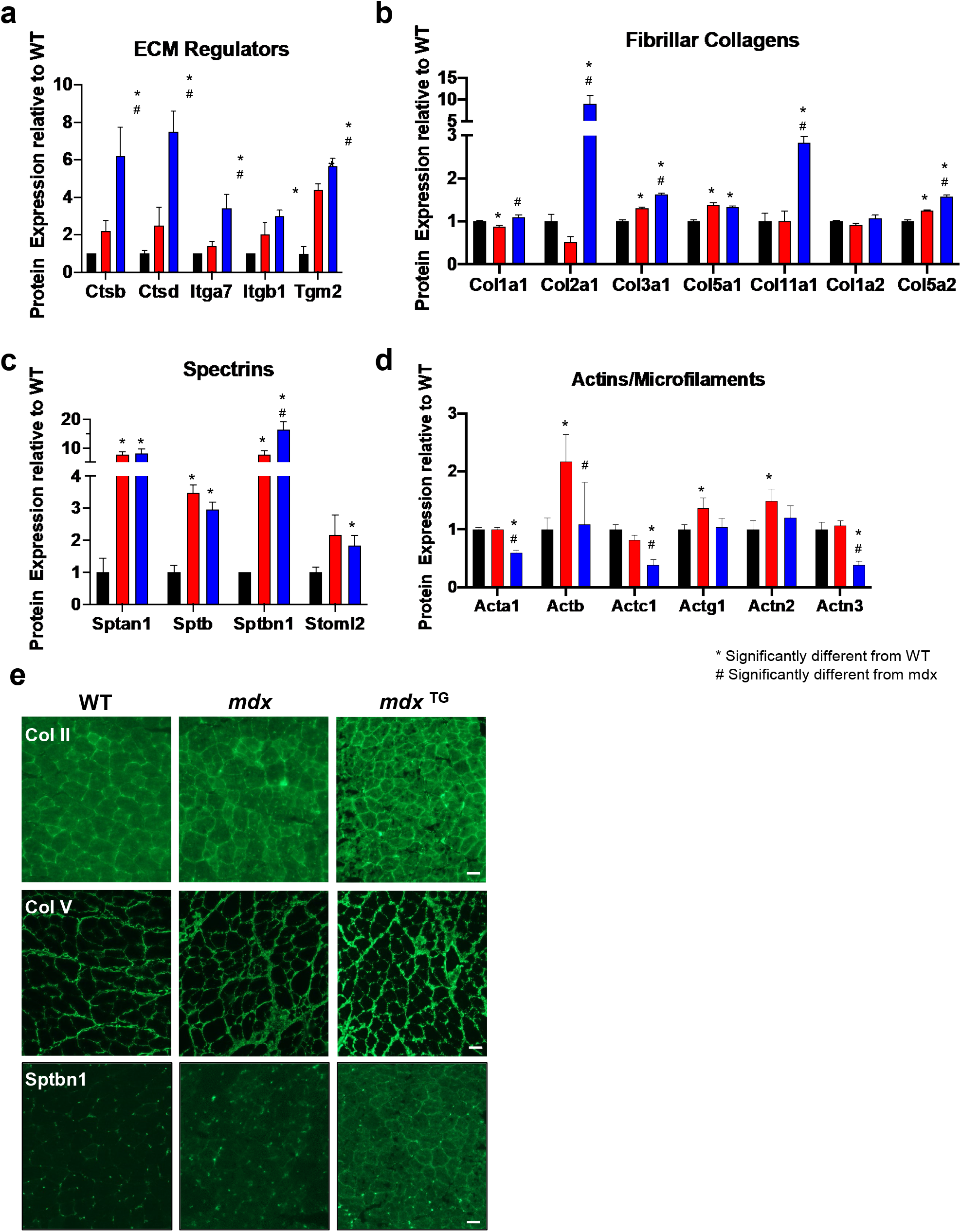
Upregulation of ECM regulators, fibrillar collagens, and spectrins in *mdx* ^TG^ muscle. Relative protein expression of ECM regulators (**a**), fibrillar collagens (**b**), spectrins (**c**) and actins/microfilaments (**d**) (*p<0.05 compared to WT, # p<0.05 compared to *mdx* by unpaired t-test). **e**) Indirect immunofluorescence of 20-week-old mouse quadricep sections using antibodies against collagen II (Col II), collagen V (Col V), and β-spectrin (Sptbn1) shows increased sarcolemmal staining in the *mdx* ^TG^ muscle compared to both WT and *mdx* muscle. Scale bar = 50 μm.

Proteomic analysis revealed compensatory upregulation of many collagens, including collagen type II (Col2) and collagen type V (Col5), along with upregulation of Sptbn1. To further interrogate the localization and abundance of these proteins, we performed indirect immunofluorescence on quadriceps sections from wild-type, *mdx*, and *mdx* ^TG^ mice with antibodies against Col2, Col5, and Sptbn1. We found increased deposition of both Col2 and Col5 at the interstitial matrix in *mdx* ^TG^ muscle (Fig. 4e), consistent with mass spectrometry results. We also observed an increase in Sptbn1 in *mdx* ^TG^ muscle at the sarcolemma compared to both *mdx* and WT and a small increase in *mdx* compared to WT, corroborating previous findings (Fig. 4e) (43). Importantly, recent findings in the golden retriever model of DMD (GRMD) revealed spectrin as a candidate protein responsible for sparing of the cranial sartorius muscle from dystrophic pathology (54). Spectrin is a mechanosensitive protein that functions as a scaffolding protein at the cell membrane to provide structural and mechanical stability (55). Additionally, spectrin is a key linker protein in the response to mechanical stimuli, coupling changes in cellular tension to downstream signaling networks, including the Hippo-YAP/TAZ network (56,57). The compensatory upregulation of spectrin along with fibrillar collagens suggests that SSPN overexpression may be rescuing the dystrophic phenotype through the alteration of mechanotransduction signaling networks.

### Ingenuity Pathway and Gene Set Enrichment Analyses Support Compensation through Mechanotransduction Pathways

To identify candidate networks and upstream regulators that drive compensatory changes in *mdx* ^TG^ muscle, we analyzed the transcriptomic and proteomic data sets using the Ingenuity Pathway Analysis (IPA) application. Based on the IPA upstream regulator algorithm and corresponding activation z-scores and p-values, we examined the top ten activated and inhibited upstream regulators for each comparison from the RNA sequencing (Fig. 5a) and mass spectrometry (Fig. 5b) data sets. Upon closer analysis of the top five activated and inhibited regulators, we observed many cytoskeletal and ECM target molecules (Table 1, Supplementary Tables 1-3, targets in bold red). Our analyses reveal that SSPN induces large-scale changes in the composition of the ECM and cytoskeleton as well as their associated signaling networks. Thus, we sought to integrate data from the transcript and protein analysis to develop a more comprehensive model reflecting the overlap of signaling networks. We performed integrative gene set enrichment analysis on the significantly differentially expressed transcripts and proteins between *mdx* and *mdx* ^TG^. This comprises the proteins and genes in the compensatory and restored categories. We observed enrichment of gene sets associated with signal transduction pathways affecting a broad spectrum of cellular and biological processes important in myogenesis, regeneration, and homeostasis of skeletal muscle involving the Ras, Rho, and Wnt signaling, actin-cytoskeleton remodeling and organization, and ECM signaling and organization (Fig. 6). These include signaling by Rho GTPases, regulation of cytoskeleton organization and cytoskeletal regulation by Rho GTPases, integrin signaling, focal adhesions formation, and response to mechanical stimulus (Fig. 6). We observed compensatory upregulation of Rho family of GTPases involved in mediating actin dynamics and actomyosin contractility including Ras homolog gene family members (RhoA and RhoC), Rac family small GTPase 1 (Rac1) and cell division cycle 42 (Cdc42). There are compensatory and restorative changes in several factors that regulate Rho GTPase activity including the GTPase activating proteins (Arhgap 6, 22, and 44), all of which are increased in *mdx* ^TG^ muscle. Wnt proteins, including Wnt4 and Wnt5a, play key roles as part of the muscle regeneration program, cytoskeletal remodeling and myoblast fusion (58,59). Wnt4 regulation of RhoA activity is important for maintaining satellite cell quiescence (59). In addition, Wnt5a and Fzd4 (rescued in *mdx* ^TG^) have been shown to regulate osteogenic differentiation after mechanical stretch (60). R-spondins family members 1 and 4 (Rspo1 and Rspo4) positively regulate the canonical Wnt signaling pathway and are important for muscle repair and regeneration (61–63). RhoA and Rac1 can also be activated by and respond to mechanical stress and stimuli. At focal adhesion sites of integrin signaling, these proteins orchestrate actin cytoskeleton remodeling important for mechanical force generation and transmission in concert with talin (Tln2) and vinculin (Vcl) (64). Several integrin and laminin subunits are also alternatively expressed in *mdx* ^TG^ relative to *mdx*. As we have shown previously, integrin alpha 7 (Itga7) is increased in *mdx* ^TG^ muscle (27). Interestingly, transcription factors Sox9 and JunB are both upregulated in *mdx* ^TG^ muscle. Sox9, important for bone and cartilage formation, is expressed in muscle progenitor cells during development and may play a role in musculoskeletal development (65). Sox9 has been shown to be a key regulator of ECM deposition, most specifically of Col2a1 (66), which is highly upregulated in *mdx* ^TG^ muscle (Fig. 4b). JunB is also important for maintaining muscle growth and hypertrophy (67). The cysteine and glycine-rich protein 3 (Csrp3, also known as muscle LIM protein, MLP) is localized at costameres and involved in promoting mechanical and stress induced myocyte differentiation and remodeling (68,69). Overall, the gene set enrichment analysis indicates that SSPN overexpression in dystrophin-deficiency results in a rewiring of signaling networks associated with cell-matrix communication and mechanotransduction, or the conversion of mechanical inputs to intracellular biochemical and biophysical signals.

**Figure 5.**
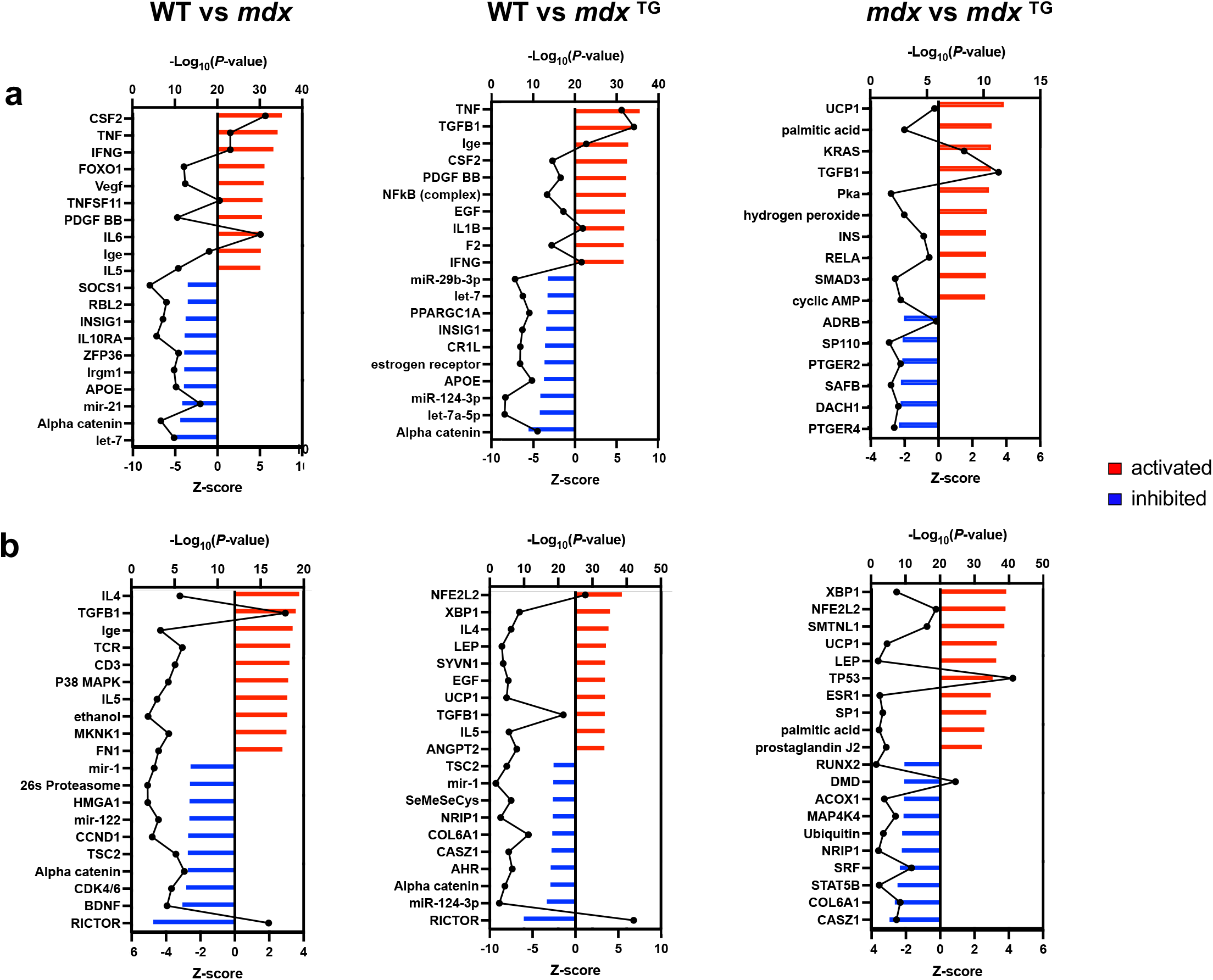
Activated and inhibited upstream regulators identified in wild-type, *mdx* and *mdx* ^TG^ muscle through Ingenuity Pathway Analysis. Ingenuity Pathway Analysis (IPA) of RNA sequencing data (**a**) and mass spectrometry data (**b**) identifying the top 10 upstream regulators that are activated (z-score, red bars) or inhibited (z-score, blue bars) with –log p-values overlaid in black. Comparisons include WT vs *mdx* (left graphs), WT vs *mdx* ^TG^ (middle graphs), and *mdx* vs *mdx* ^TG^ (right graphs).

**Figure 6.**
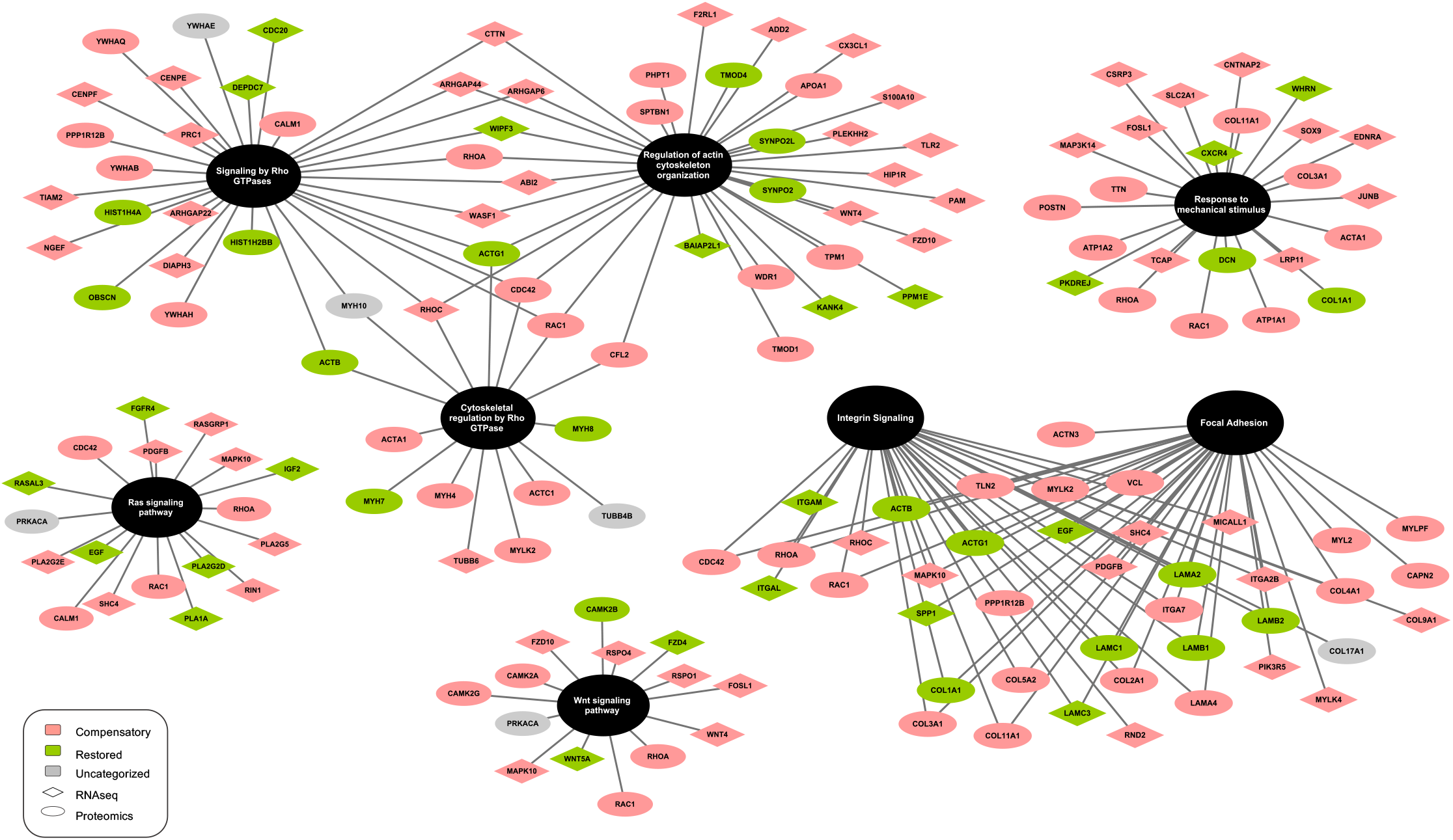
Integrated gene set enrichment analysis identifies differential expression of mechanosignaling pathways in *mdx* ^TG^. To develop an integrated model of overlapping signaling networks driving rescue of the *mdx* ^TG^ skeletal muscle, we combined transcripts (diamonds) and proteins (ovals) that were differentially expressed between *mdx* and *mdx* ^TG^ tissue and performed gene set enrichment analysis. Compensatory changes are in pink, restored changes are in green, and uncategorized changes are in gray. The gene set enrichment analysis highlights changes in Rho, Rac, Wnt and integrin signaling in addition to cell adhesion and response to mechanical stimulus.

**Table 1.**
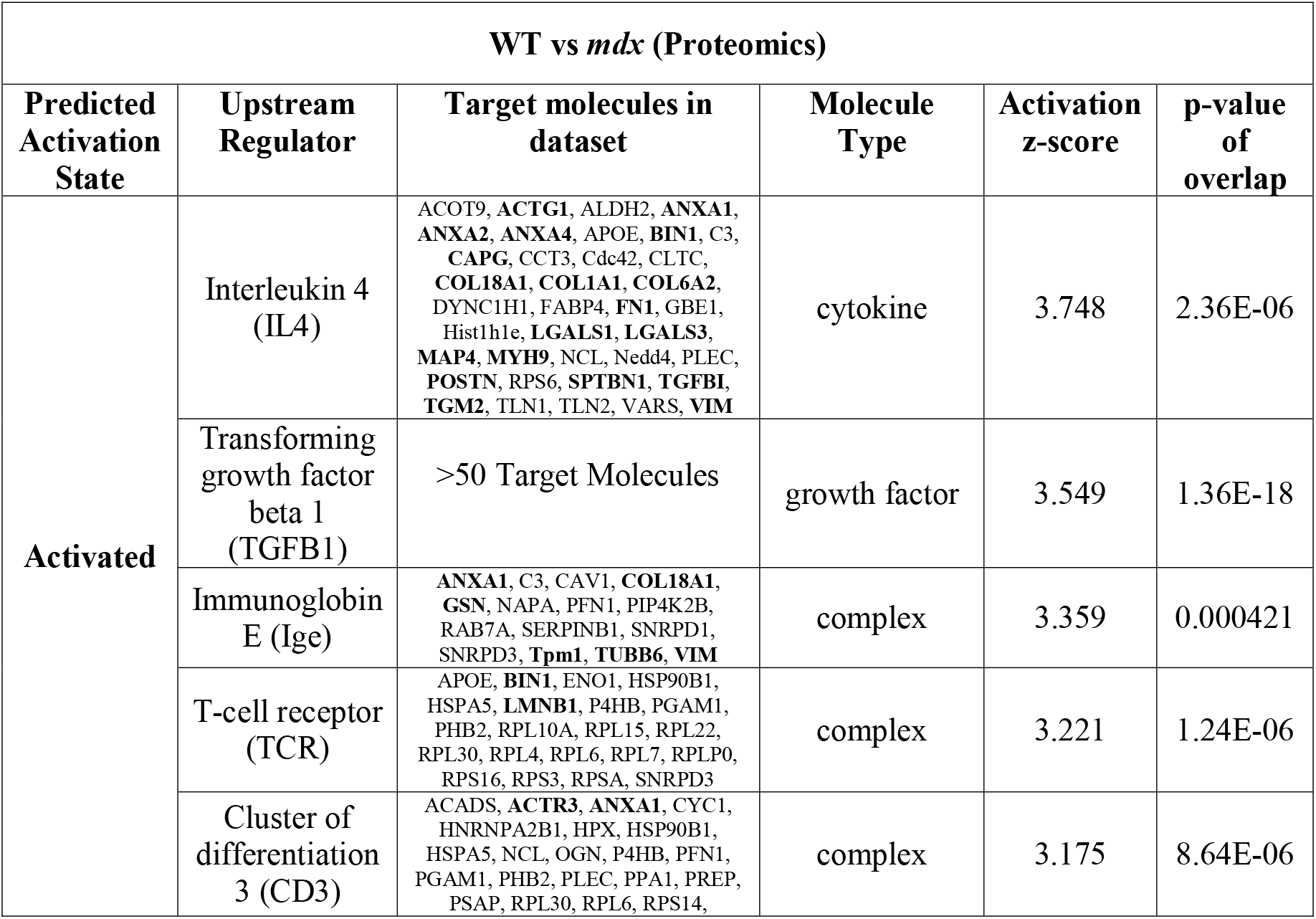
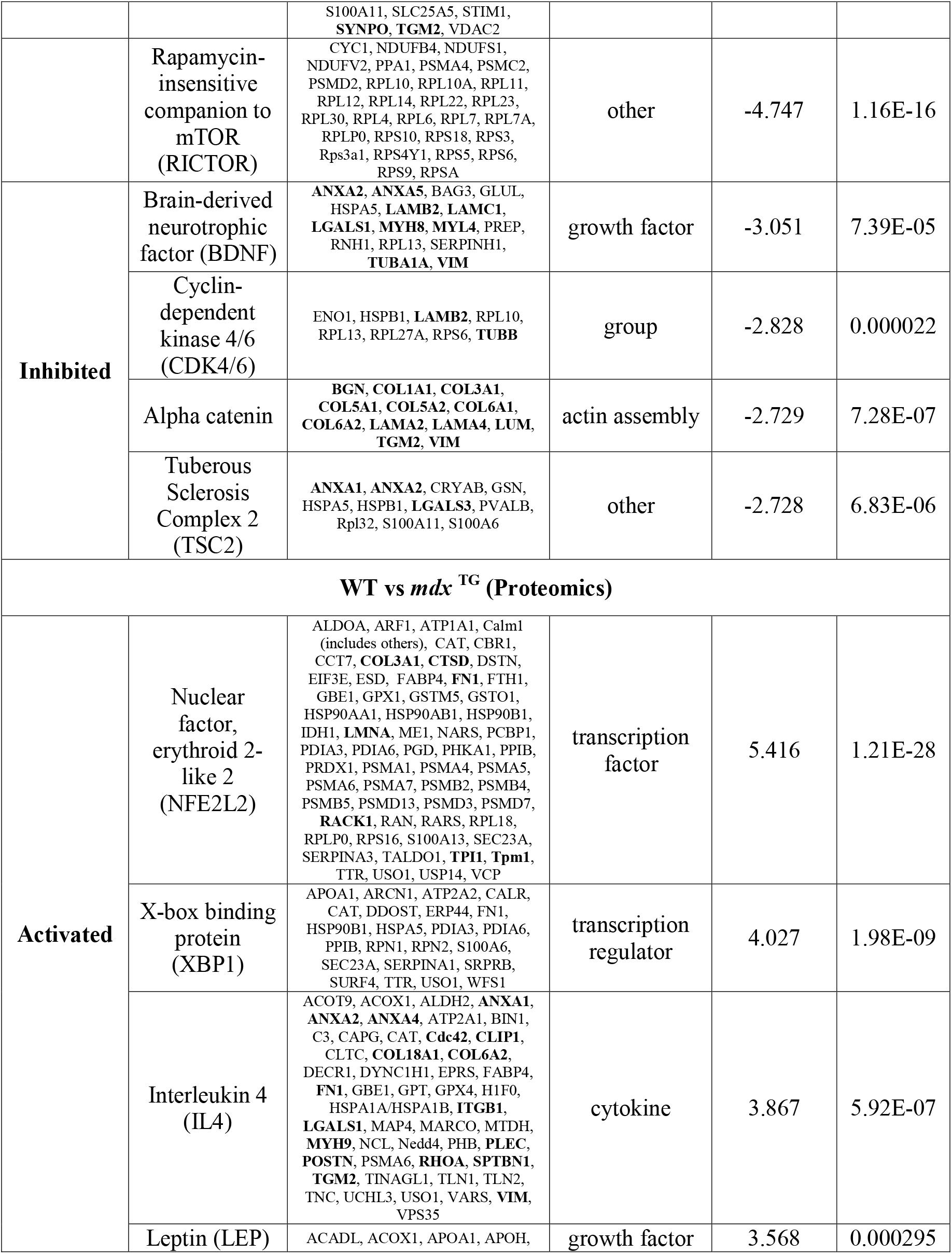
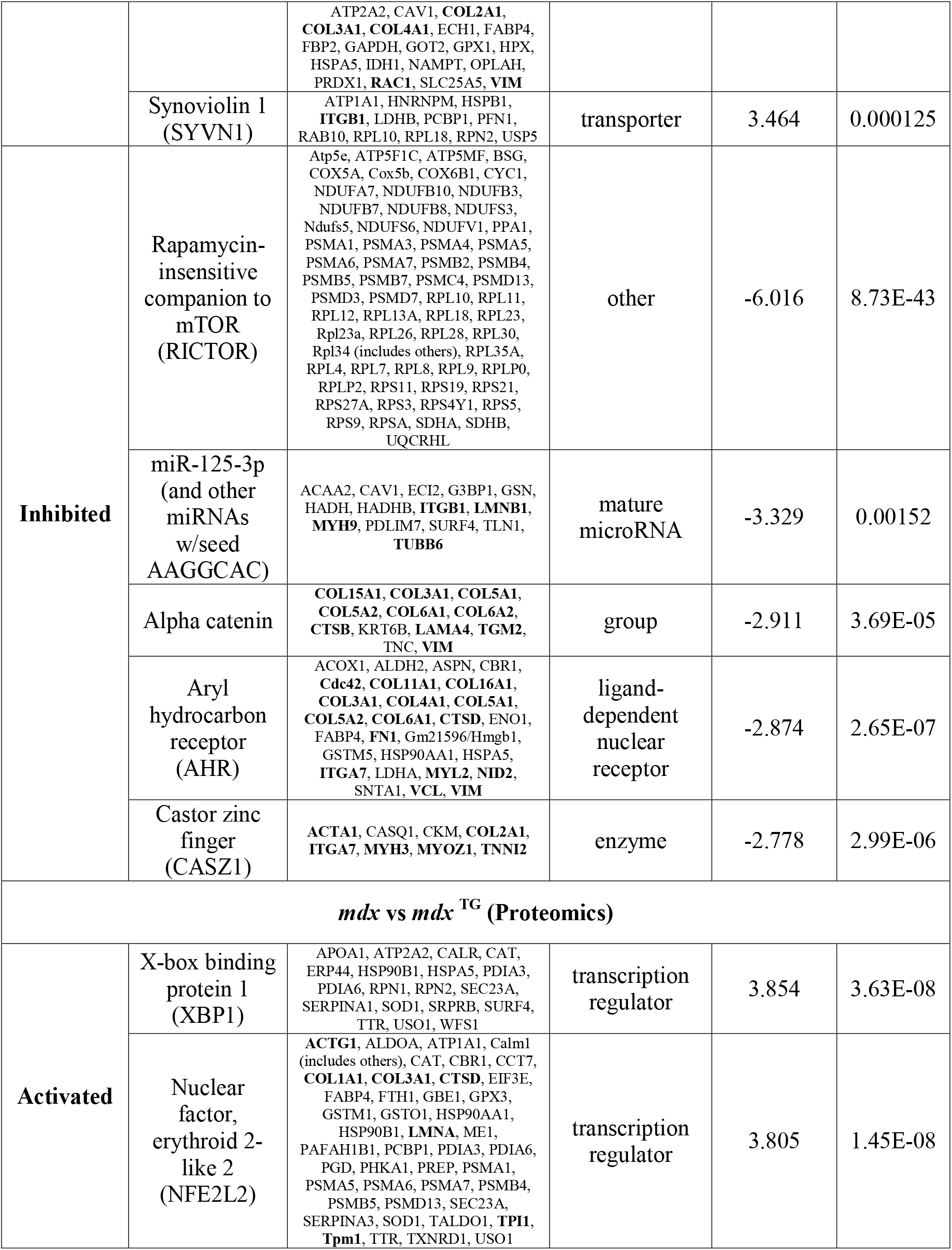
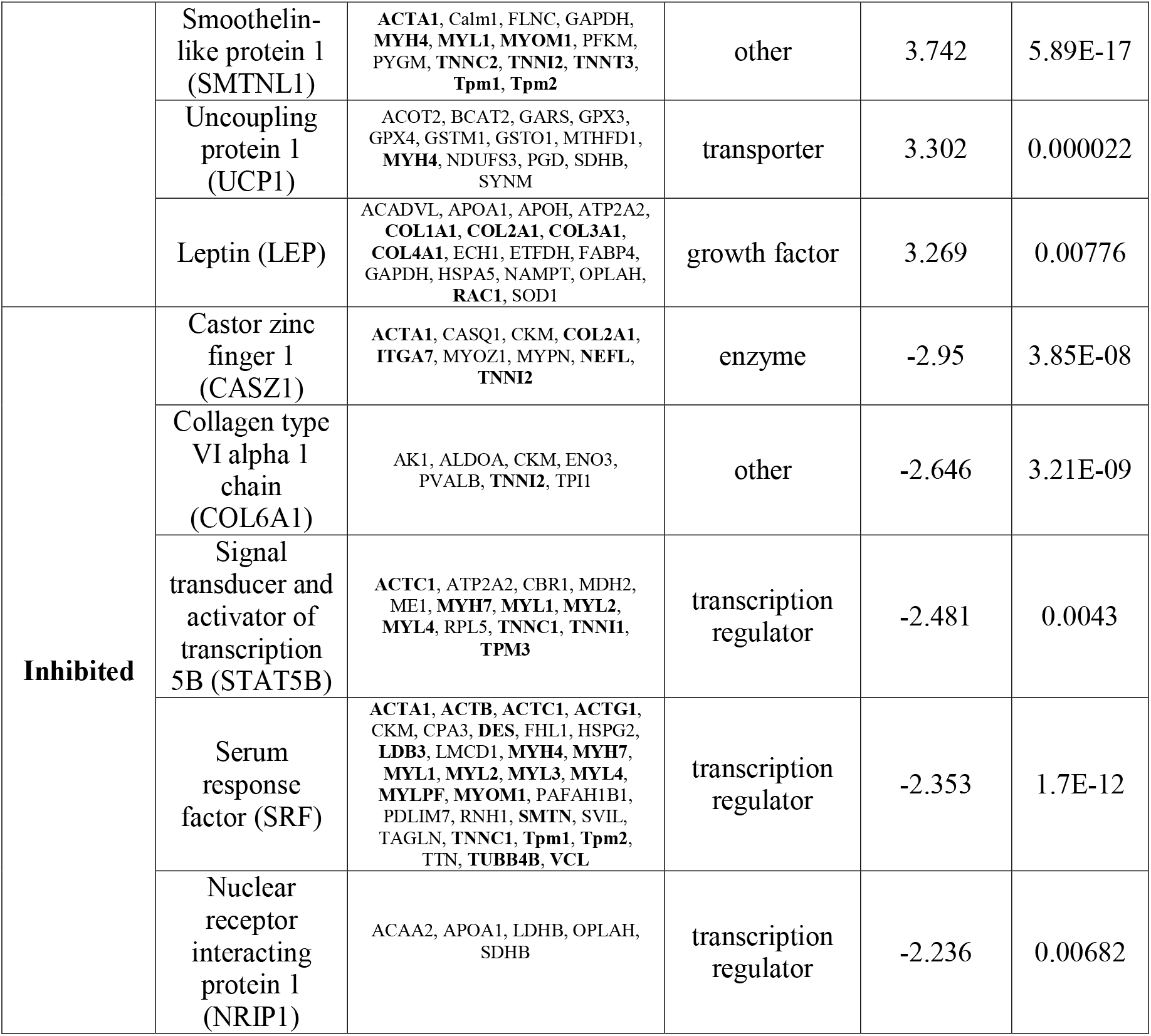
Ingenuity Pathway Analysis Summary – Upstream Regulators.

## DISCUSSION

In our multi-omics approach to identifying mechanisms of SSPN amelioration of dystrophic pathology, we built on previous comparative transcriptomic and proteomic studies that have identified the molecular changes in *mdx* muscle throughout progression of disease in muscle groups that are differentially affected by disease (37,40,42,70). Our approach combines RNA sequencing transcriptomics with ECM-optimized capture techniques to identify SSPN-induced cellular pathways. Findings from our proteomic analysis reveal compensatory alterations in the abundance of collagens in *mdx* ^TG^ muscle, especially the highly glycosylated collagens II, V, and XI. Collagen is the most abundant protein in the ECM and its high tensile strength allows transmission of forces generated by skeletal muscle fibers. The most abundant collagens in skeletal muscle are collagens I, III, IV, and VI. While collagens were upregulated in the *mdx* ^TG^, they were primarily collagens not typically observed in skeletal muscle, or those of low abundance, including collagens II, V, and XI (71). Collagen II is typically found in articular cartilage and intervertebral discs and is highly glycosylated (72,73). The presence of these bulky disaccharide groups is believed to hinder the formation of highly ordered fibrils, and has been shown to increase substrate compliance in co-cultures with collagen I (74). Collagens V and XI are typically observed only during skeletal muscle development, however, the function of collagen XI in skeletal muscle is unknown. Baghdadi and colleagues (75) recently reported that satellite cells produce collagen V that is critical for the calcitonin receptor and notch signaling cascade, which maintains satellite cells in a quiescent state. Taken together, the changes observed in the *mdx* ^TG^ ECM suggest that upregulation of these developmental and cartilaginous collagens may be beneficial to muscle.

The identification of β-spectrin upregulation as another potential compensatory protein is supported by data from the GRMD dog model of DMD. Nghiem and colleagues (54) identified spectrin as a candidate protein in a study of the cranial sartorius muscle that is spared from dystrophic pathology through compensatory hypertrophy *via* myostatin signaling. Spectrin is a scaffolding protein that functions to localize and stabilize surface proteins in nonerythroid cells, anchoring them to the cytoskeletal actin network (76). The repeated triple helical units of the spectrin rod domain would support this function by stabilizing surface proteins and mediating cell-matrix and cell-cell interactions. Interestingly, dystrophin contains similar spectrin-like repeats in its central rod domain and has been reported to bind spectrin (77–79). Data from the IPA and gene enrichment analysis also identify spectrin as a target molecule of signaling pathways associated with a reorganization of the actin cytoskeleton. Together, the upregulation of collagens and scaffolding cytoskeletal components such as spectrin, suggest that SSPN overexpression and the increase of adhesion complexes at the membrane may induce a rewiring of signaling between the ECM and the cytoskeleton to stabilize the sarcolemma during contraction and enhance mechanotransduction in order to improve force transmission.

Mechanotransduction has been described in the *mdx* mouse model in many studies that highlight the role of microtubules, adhesion complexes, mechanosensitive ion channels, and YAP/TAZ signaling in both skeletal and cardiac muscle pathology (80–86). Our study corroborates some of these findings when comparing wild-type and *mdx* muscle (Supplemental Fig. 2). In comparing wild-type and *mdx* to our *mdx* ^TG^ model; however, we do not observe a restoration of these pathways, suggesting that SSPN is improving force transmission and muscle function through alternative pathways. Our gene set enrichment analysis revealed potential candidate alternative pathways involved in mechanotransduction through regulation of cytoskeleton, focal adhesion, and integrin signaling (Fig. 6). Notably, we identified compensatory changes in RhoA, Rac, and Wnt signaling molecules in the *mdx* ^TG^ muscle. Reciprocal crosstalk between the Wnt, Rho GTPase and the integrin complex mediate cell adhesion signaling cascades and regulation of the cytoskeleton (87–89). Integrins are bidirectional mechano-transducers that detect mechanical cues and translate these signals to affect intracellular and extracellular behavior and response. Through its interaction with adaptor proteins at sites of focal adhesion, the integrin complex regulates the dynamic, cyclic, spatial and temporal activation of RhoA, Rac1 or Cdc42 leading to actin polymerization and depolymerization which drives force propagation, cell motility and contractility (87,90,91). RhoA, through downstream effectors, stabilizes actin filaments and regulates the activity of cofilin 2 (CFL2) which together with WD repeat-containing protein 1 (Wdr1) engages in depolymerization and severing of actin filaments, important to replenishing the actin monomer pool. While Rac1 and cdc42 are involved with activation of Wiskott-Aldrich syndrome protein family member 1 (Wasf1) and Abl interactor 2 (ABI2), members of the WAVE complex, to promote actin nucleation and branching (92). Obscurin (Obscn), a sarcomeric rho-guanine nucleotide exchange factor (93), has been shown to activate RhoA in skeletal muscle (94). Adducin 2 (Add 2) functions to cap the barbed end of actin filaments and is involved with recruiting spectrin tetramers to actin filaments (95,96). Rac1 and cdc42 have also been reported to signal through the DGC complex and may be downregulated during muscle atrophy (97). Furthermore, RhoA, in addition to Rac1 and Cdc42, regulate transcription by SRF (98). Interestingly, the transcription factor SRF is in one of the top five significantly inhibited/affected upstream regulators in our IPA analysis between *mdx* and *mdx* ^TG^ (Table 1).

## CONCLUSIONS

We generated a data-driven schematic of the *mdx* muscle rescued by SSPN overexpression that highlights compensatory changes in the ECM and cytoskeleton at both the transcript and protein level (Fig. 7). Our multi-omics data suggest that SSPN rescues dystrophin deficiency partially through a rewiring of cell-matrix interactions that may enhance mechanotransduction signaling cascades and improve lateral force transmission.

**Figure 7.**
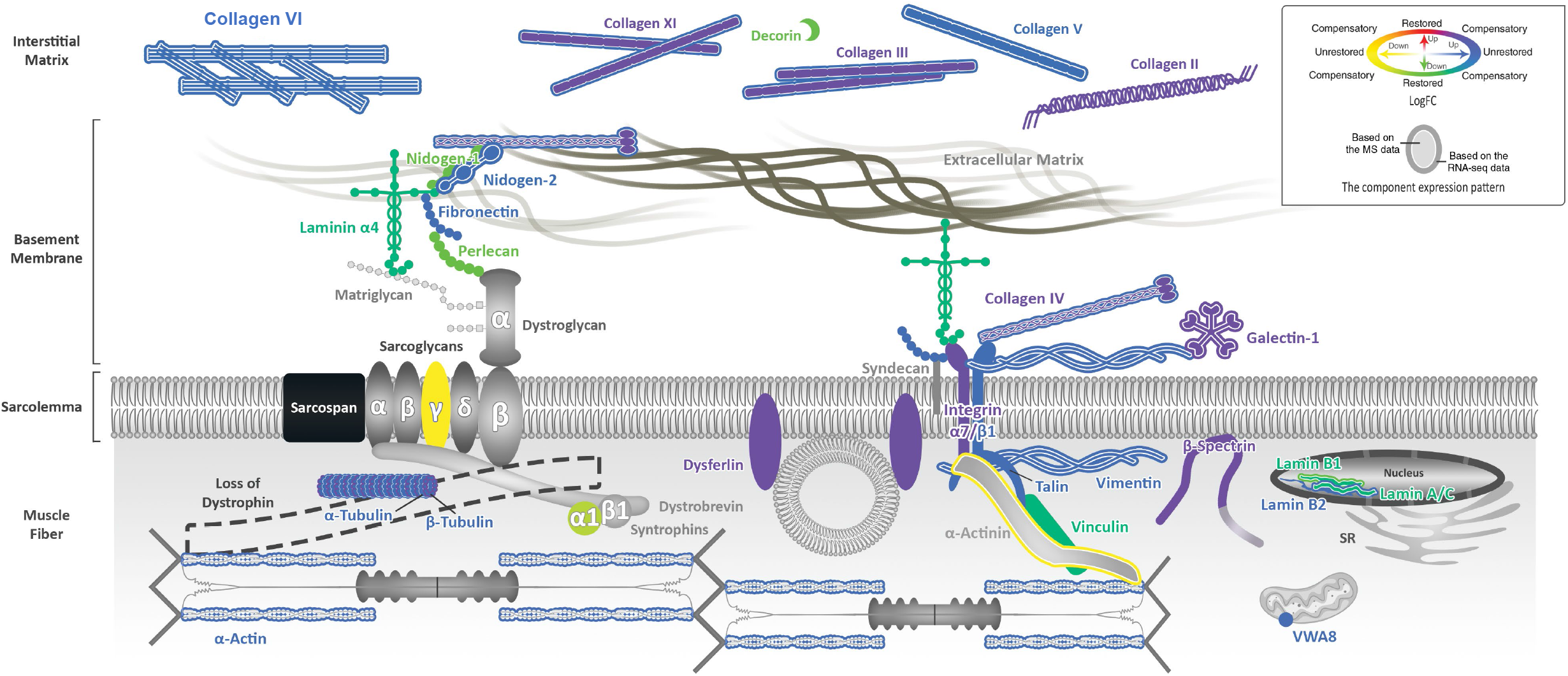
SSPN overexpression rewires ECM-cell communication. Schematic summarizing ECM and cytoskeletal changes in RNA sequencing and proteomic data sets. Changes in gene or protein expression are represented by color according to the key, indicating compensatory upregulation of many ECM and cytoskeletal molecules with downstream signaling effects (in purple). The x and y axes in the color key correspond to *mdx* ^TG^ relative to WT and *mdx* ^TG^ relative to *mdx*, respectively.

## LIST OF ABBREVIATIONS

DGC: dystrophin-glycoprotein complex
ECM: extracellular matrix
SSPN: sarcospan
DMD: Duchenne muscular dystrophy
SG: sarcoglycans
hSSPN: human sarcospan
mSSPN: mouse sarcospan
CPM: counts per million
DEG: differentially expressed genes
BSA: bovine serum albumin
PBS: phosphate buffered saline
GO: gene ontology
GRMD: golden retriever model of DMD
IPA: Ingenuity Pathway Analysis
ESR1: estrogen receptor 1
Col2: collagen type II
Col5: collagen type V
Sptbn1: non-erythrocyte β-spectrin
Rac1: Rac family small GTPase 1
Cdc42: cell division cycle 42
Arhgap: GTPase activating proteins
TLN2: talin
VCL: vinculin
Itga7: integrin alpha 7
Wasf1: Wiskott-Aldrich syndrome protein family member 1
ABI2: Abl interactor 2
Obscn: Obscurin
Add 2: Adducin 2
CFL2: cofilin 2
Wdr1: WD repeat-containing protein 1
Csrp3: cysteine and glycine-rich protein 3
Sox9: SRY-Box Transcription Factor 9
JunB: JunB proto-oncogene, AP-1 transcription factor subunit
YAP: yes-associated protein
TAZ: transcriptional coactivator with PDZ-binding motif
Col2a1: collagen type II alpha 1 chain
Rho: Ras homolog
RAC: Rac family small GTPase
Wnt4: wingless-type MMTV integration site family, member 4
Wnt5: wingless-type MMTV integration site family, member 5
NAD/NADP: Nicotinamide adenine dinucleotide/Nicotinamide adenine dinucleotide phosphate
GTPases: guanosine triphosphate hydrolases
Ras: Rat sarcoma virus family protein
RhoA: Ras homolog family member A
RhoC: Ras homolog family member C
Fzd4: frizzled class receptor 4
Rspo1: R-spondin 1
Rspo4: R-spondin 4
cdc42: cell division cycle 42
WAVE complex: WASP family Verprolin homolog complex

## DECLARATIONS

### Ethics approval and consent to participate

Not Applicable

### Consent for publication

Not Applicable

### Availability of data and materials

The proteomics dataset supporting the conclusions in this article is available in the PRIDE Archive repository. The RNA sequencing dataset supporting the conclusions in this article will be available on the NCBI GEO database upon publication.

### Authors’ Contributions

CS, EG, JLM, KSR, HM, and RHC conceptualized the project, study design, and methodology. RHC oversaw and coordinated the project. YZK performed initial computational analysis of the RNA sequencing data and JLM, HM, CS, EG, and KSR further developed the analysis, visualization, and interpretation of the results. KH and LRS performed the mass spectrometry with initial analysis and KSR, JLM, EG, and HM further developed analysis and interpreted the results. CS, JLM, KSR, EG analyzed and interpreted the Ingenuity Pathway Analysis. MHA conceptualized and performed the gene ontology enrichment analysis and data visualization. KMS validated key findings using immunofluorescence microscopy. HM proposed the rewiring concept and performed the integrative gene set enrichment to develop the model. JLM, KSR, and HM drafted the main manuscript and JLM, KSR, HM, CS, EG, MHA prepared the figures in consultation with RHC. All authors reviewed and approved the final manuscript.

## Acknowledgements

We thank Dr. Ekaterina Mokhonova (UCLA) and Katherine Hammond for providing all tissues and mouse colony oversight, genotyping, and maintenance. This work was supported by the QCB Collaboratory community directed by Dr. Matteo Pellegrini (UCLA).

