## Supplemental Figures for "Multi-omics analysis of sarcospan overexpression in *mdx* skeletal muscle reveals compensatory remodeling of cytoskeleton-matrix interactions that promote mechanotransduction pathways"

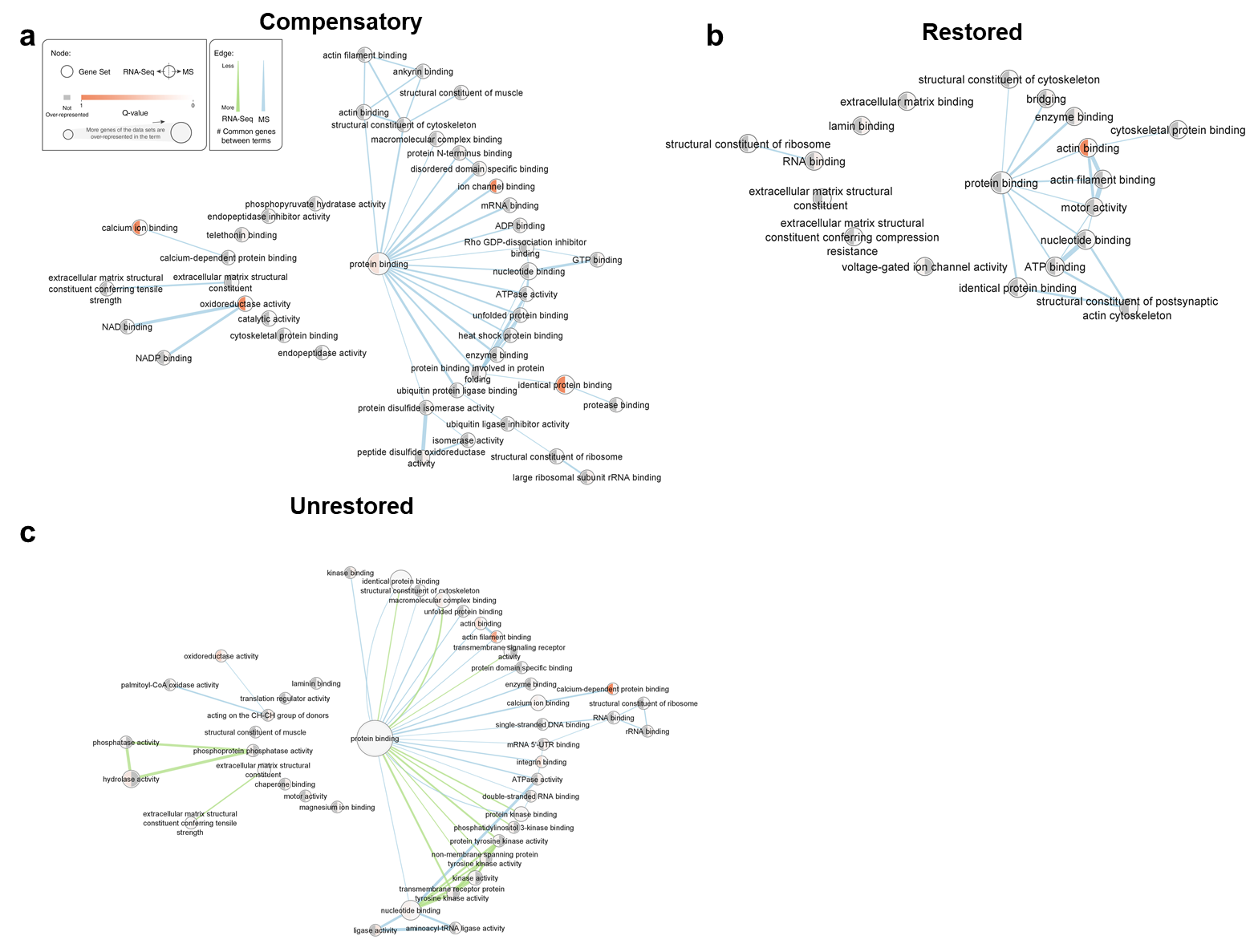

**Supplemental Figure 1.** Enrichment analysis was performed using the Database for Annotation, Visualization and Integrated Discovery platform (DAVID, version 2021) and identified Gene ontology terms, denoted as nodes for both the RNA sequencing (right half of nodes) and mass spectrometry (MS) datasets (left half of nodes). The level of enrichment is based on the Q-value and is indicated by a red color spectrum. The thickness of the connecting lines in green (RNAseq) or blue (MS) indicates the # of common genes between the connecting terms. The enrichment analysis was performed on gene or protein lists from compensatory (**a**), restored (**b**), or unrestored (**c**), categories.

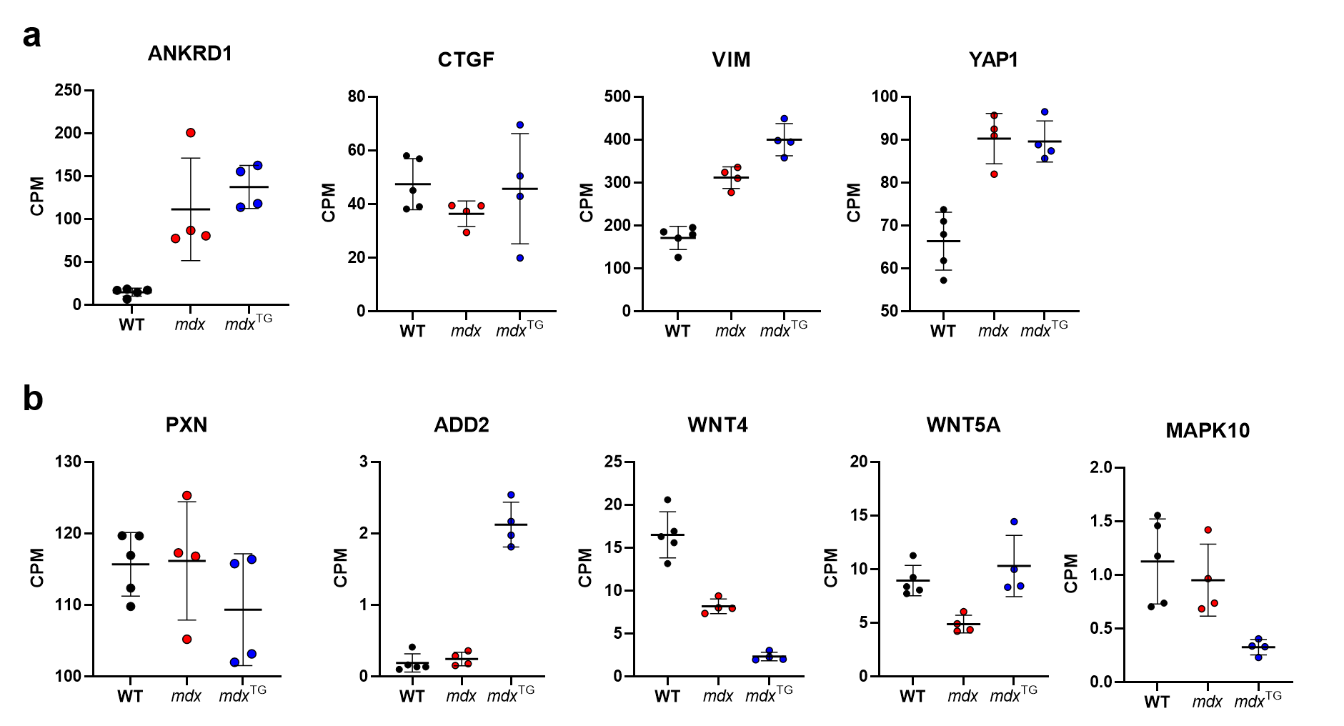

**Supplemental Figure 2.** Gene expression from RNA sequencing of genes associated with YAP/TAZ signaling (a) or Wnt signaling (b) in WT, *mdx*, and *mdx* ^TG^ muscle in counts per million (CPM). ANKRD1: Ankyrin Repeat Domain 1; CTGF: connective tissue growth factor; VIM: vimentin; YAP1: yes associated protein 1; PXN: paxillin; ADD2: adducin 2; WNT4: wingless-type MMTV integration site family, member 4; WNT5A: wingless-type MMTV integration site family, member 5a; MAPK10: mitogen-activated protein kinase 10.

Supplementary Table 1. Ingenuity Pathway Analysis – RNAsequencing WT vs *mdx*

*Target genes in red are ECM, ECM-associated, cytoskeletal, or cytoskeletal-associated genes.

| **Predicted Activation State** | **Upstream Regulator** | **Molecule Type** | **Target Genes** | **Activation z-score** | **p-value of overlap** |
| --- | --- | --- | --- | --- | --- |
| **Inhibited** | let-7 | microRNA | AURKB, BCAT1, BRCA1, BUB1, BUB1B, CASP3, CCNA2, CCNB1, CCNF, CD44, CDC20, CDCA2, CDCA3, CDCA7, CDCA8, CDK1, CDK6, CDT1, **COL3A1**, **DMD**, E2F8, FANCD2, FOSL1, ID2, IGF2BP2, MCM5, NUF2, PLAGL1, PLAUR, RRM2, SKP2, SOX9 | -4.845 | 1.42E-10 |
|  | Alpha catenin | actin assembly | **ADAM8**, **ADAMTS4**, BCL3, C1QTNF3, **COL15A1**, **COL3A1**, **COL6A3**, EPHA3, IGF2 , IL2RG, **ITGAM**, LUM, LYZ, **MMP12**, **MMP19**, SOCS3, **TIMP1**, TNFAIP2, TNFRSF12A, VCAM1 | -4.406 | 1.85E-07 |
|  | mir-21 | microRNA | AIF1, ANLN, ASPM, BCL2, CCNA2, CCNB1, CD180, CDK6, CDKN1A, CDKN2A, CLEC5A, COL3A1, CSF3R, DDAH1, DLGAP5, ECT2, FCGR1A, FCGR2A, FGL2, IFI16, IGHM, INPP5D, IRGM, KIF23, KIF4A, KIFC1, KNTC1, MKI67, NPAS2, NUSAP1, OAS3, PBK, PER2, PRC1, RAB7B, RACGAP1, RAD51AP1, Slfn2, SMC2, STMN1, **TGFB1**, TIAM1, TLR1, TOP2A, TRAF1, TREM2, VCAM1, ZWILCH | -4.166 | 9.62E-17 |
|  | Apolipoprotein E (APOE) | transporter | ADGRE1, APOE, ATF3, C5AR2, CASP1, CASP3, Ccl2, CCL22, CCR5, CD44, CD68, CTSS, CYBB, DAGLA, EGR1, FCGR1A, FCGR2B, FOS, IL10RA, IL1RN, **ITGAM**, **ITGAX**, LIPA, LPL, LRP8, **MMP3**, MSR1, NCF1, NCF2, NPNT, PRKCB, SERPINA3, SOCS3, SPP1, TCIRG1, TGFB1, TIAM1, TIFA, **TIMP1**, TNIP1, TREM2, UCP2, VCAM1 | -3.966 | 4.89E-11 |
|  | immunity-related GTPase family M member 1  (IRGM1) | GTPase | AURKB, BUB1, Ccl2, CCNA2, CCNB1, CCNB2, CDCA3, DTL, ID2, IFI16, KIF20A, LILRB4, MKI67, NCAPG, NEK2, RRM2 | -3.93 | 1.45E-10 |
| **Activated** | Vascular endothelial growth factor (VEGF) | growth factor | ALOX5AP, ANPEP, ARHGAP22, ASNS, ATF3, AURKB, BCL2, BIRC5, BUB1, BUB1B, CASP1, CASP3, Ccl2, CCNF, CD44, CDC20, CDK1, CDKN2A, CENPF, CLDN1, CNTFR, CXCR4, DUSP5, EGR1, EMP1, FOSL1, FOXM1, GJB2, GPRC5B, GPSM2, GRB10, HELLS, HPSE, KIF11, KIF15, KIF20B, KIF22, LRP8, LY75, MCM5, MELK, MKI67, **MMP14**, Mt2, NEK2, NRCAM, PLAUR, **PLK1**, **PLK4**, PRC1, PRKCB, SCML4, SFN, SKP2, SMC2, SOCS3, TGFB1, TIMP1, TNFRSF11A, VAV3, VCAM1 | 5.457 | 3.46E-13 |
|  | Forkhead Box O1 (FOXO1) | transcription regulator | ANLN, ASPM, ATP6V0D2, BCL2, BID, BIRC5, C1QA, CCNA2, CCNB1, CCNB2, CCNF, CDK1, CDKN1A, CDKN2A, CENPF, CIDEA, DCX, DIO2, DLGAP5, EGR1, FABP5, FOS, FOXM1, IKZF1, IL17RA  IL7R, **ITGAM**, **ITGB2**, KIF11, LCP2, LPL, MCM5, **MMP3**, **MYL3**, MYOG, NCAPG, NEK2, NUSAP1, PAK1, PBK, PRC1, PTPRC, Ptprv, RUNX2, SFN, SLC25A22, SLC25A24, SPC25, SQLE, **TGFB1**, TNFRSF11A, TRAF1, TSPO, VCAM1, WNT4 | 5.554 | 7E-13 |
|  | Interferon Gamma (IFNG) | cytokine | ADCY7, ADRA2A, AIF1, AIF1L, ALOX5AP, ASNS, ATF3, BCL2, BCL3, BID, BMP6, BST1, C1QA, C1QB, C1QC, CASP1, CASP3, Ccl2, CCL22, CCL28, Ccl7, Ccl8, CCNA2, CCR1, CCR5, CD14, CD276, CD4, CD44, CD55, CD68, CD72, CD83, CDK5R1, CDKN1A, CDKN2A, CELSR2, CERS6, CHRND, CHRNG, CIITA, CLEC5A, CLEC7A, CORO1A , CSF2RB, **CTSH**, **CTSS**, CX3CR1, CXCL16, CXCR4, CYBA, CYBB, DUSP5, E2F1, EFCAB6, EGR1, FABP5, FCER1G, FCGR1A, FCGR2A, FCGR2B., FCGR3A/FCGR3B, FGL2, FOS, GDF15, GJB2, GPRC5B, HCK, HLA-DMB, IFI16, IFI30, IKBKE, IL10RA, IL17RA, IL18RAP, IL1RN, IL7R, IRF8, IRGM, **ITGAL**, **ITGAM**, **ITGAX**, **ITGB2**, CNMA1, KLF10, **LAMC2**, LAT2, LCP2, LGALS3, LPL, MEFV, MERTK, **MMP12**, **MMP3**, Ms4a4b (includes others), MSR1, Mt1, MYOG, NAPSA, NCAM1, NCF2, NEURL3, OAS3, P2RY6, PARVG, PBK, PLA2G5, PLAUR, PLEK, PTAFR, PTPN6, RAC2, RUNX2, RUNX3, SBNO2, SGPL1, SLC11A1, SLC15A3, SOAT1, SOCS3, SPI1, **SPP1**, SPRR1A, SQLE, STX11, TBXAS1, TCIRG1, TFRC, **TGFB1**, THEMIS2, **TIMP1**, TLR1, TLR6, TLR7, TLR8, TNFAIP2, TNFRSF11A, TNFRSF12A, TNFRSF1B, TREM2, TYROBP, VCAM1, Wfdc17 | 6.595 | 8.6E-24 |
|  | Tumor necrosis factor (TNF) | cytokine | AATK, ABCC3, **ADAM8**, **ADAMTS4**, AKR1B10, ALCAM, ALOX5AP, ANPEP, APOE, ARHGAP22, ATF3, B4GALNT1, BCL2, BCL3, BID, BIRC5, BUB1B, C3AR1, CASP1, CASP3, Ccl2, CCL22, CCL28, Ccl7, CCNE1, CCR1, CCR5, CD14, CD247, CD4, CD44, CD55, CD83, CDK5R1, CDKN1A, CDKN2A, CERS6, CHRND, CHRNG, CIITA, CLEC5A, CMBL, CNR2, **COL15A1**, **COL3A1**, COTL1, CSF2RB, **CTSS**, CX3CR1, CXCL16, CXCR4, CYBA, CYBB, CYP1B1, CYTIP, **DMD**, DUSP14, DUSP5, E2F1, EFHD2, EGR1, ELF3, EMP1, EXOC3L4, FABP5, FCER1G, FCGR2B, FOS, FOSL1, FRZB, FUT4, GABRA1, GDF15, GPR176, GPRC5B, GRN, H19, HEXB, HGF, HK3, Hmgn2 (includes others), HPGDS, IDH2, IFI16, IGF2, IKBKE, IL10RA, IL1RN, IL21R, IL7R, INPP5D, IRF8, **ITGAL**, **ITGAM**, **ITGAX**, **ITGB2**, KCNH2, KIF20A, KLF10, KLF5, KRT18, **LAMC2**, **LGALS3**, LPL, MBP, MEFV, **MMP12**, **MMP14**, **MMP3**, MSLN, MSR1, MSTN, Mt1, Mt2, MYOG, NAIP, NCAM1, NCF1, NCF2, OAS3, P2RY6, PCDH7, PDK3, PDPN, PER2, PLA2G5, PLAUR, PLD3, **POSTN**, PTPRC, PYCARD, RAB32, RASSF7, RGS1, RGS16, RRAD, RRM2, RUNX2, SCUBE2, SELPLG, SERPINA3, SERPINB1, SGPL1, SLC11A1, SLC15A3, SLC22A4, Slfn2, SOAT1, SOCS3, SOX9, **SPP1**, SQLE, STMN1, TBXAS1, TFRC, **TGFB1**, TIFA, **TIMP1**, TLR7, TLR8, TNFAIP2, TNFRSF11A, TNFRSF1B, TNIP1, TRAF1, TREM2, UCP2, VAV3, VCAM1, WISP1, ZNF365, ZNF750 | 7.103 | 8.45E-24 |
|  | Colony stimulating factor 2 (CSF2) | cytokine | ABCG1, ADAl, **ADAM8**, ALOX5AP, ANLN, BCL2, BCL3, BID, BIRC5, BUB1, BUB1B, CASP1, CASP3, Ccl2, Ccl8, CCNA2, CCNB1, CCNF, CCR1, CCR5  CD14, CD180, CD276, CD83, CDC20. CDCA2, CDCA3, CDCA8, CDK1, CDKN1A, CENPE, CIITA, CLEC6A, CLEC7A, **COL8A1**, CSF2RA, CSF2RB, CXCR4, CYBB, E2F8, EGR1, FANCA, FCGR1A, FCGR2B, FIGNL1, FOS, FOSL1, FOXM1, GDF15, HGF, ID2, IL1RN, INPP5D, **ITGAM**, **ITGAX**, KIF11, KNTC1, LCP1, LY75, LY9, MCM5, MKI67, **MMP14**, MNS1, NEK2, NUSAP1, PLK1, PRC1, PTGER2, PTK2B, RACGAP1, **RHOH**, RRM2, SLAMF7, Slfn2, SMC2, SNTB1, SOCS3, SPC25, SPI1, **SPP1**, STMN1, **TGFB1**, TIFA, TLR1, TNFRSF11A, TNFRSF1B, TOP2A, UHRF1 | 7.612 | 4.74E-32 |

Supplementary Table 2. Ingenuity Pathway Analysis – RNAsequencing WT vs *mdx* ^TG^

*Target genes in red are ECM, ECM-associated, cytoskeletal, or cytoskeletal-associated genes.

| **Predicted Activation State** | **Upstream Regulator** | **Molecule Type** | **Target Genes** | **Activation z-score** | **p-value of overlap** |
| --- | --- | --- | --- | --- | --- |
| **Inhibited** | Alpha catenin | actin assembly | **ADAM8**, **ADAMTS12**, **ADAMTS4,** BCL3, BGN  BIRC3, CDH11, **COL15A1**, **COL3A1**, **COL5A1**, **COL5A2**, **COL6A1,** **COL6A2**, **COL6A3**, **CTSB**, CXCL10, EPHA3, IGFBP4, **ITGA5**, KLK3, **LUM**, LYZ, **MMP12**, **MMP19**, **MMP2**, PTGS2, RELB, **RHOC**, **TIMP1**, **TNC**, TNFAIP3, TNFRSF12A, **VIM** | -5.581 | 8.36E-12 |
|  | let-7a-5p (and other miRNAs w/seed GAGGUAG) | mature microRNA | AURKB, BIRC5, CAPG, CASP3, CCNE1, CDK6, CDKN2A, **COL27A1**, **COL3A1**, DOCK5, FADS2, HMGA1, Hmga2, IGF2BP2, PTGS2, **RHOG**, S100A4, SLC25A24, SMOX, THBS1, TLR4, UHRF1, **VIM** | -4.24 | 0.00064 |
|  | miR-124-3p (and other miRNAs w/seed AAGGCAC) | mature microRNA | ANXA8/ANXA8L1, CDK6, CHSY1, EGR1, ELOVL1, FAM129B, FAM83H, GAS2L1, HTATIP2, INO80C, KLF15, LDLR, LITAF, MDFIC, NME4, OAF, PGF, PGM1, PLP2, RASSF5, RBM47, **RHOG**, SERPINB6, SOX9, STOM, SUCLG2, TMBIM1, TSC22D4, **TUBB6**, UHRF1, ZFP36L2 | -4.172 | 0.000432 |
|  | Apolipoprotein E (APOE) | transporter | ACAT1, ADGRE1, ALOX12, ATF3, BEGAIN, BGN, CASP3, CCL22, CCL5, CCR5, CD44, CD68, CD80, CD86, CLU, **COL18A1**, CREB3L1, **CTSB**, **CTSK**, **CTSS**, Cyb5r3, CYBB, EGR1, EMILIN1, F2R, F2RL1, FOS, GLUL, GRIA3, HSPA1A/HSPA1B, Hspa1b, HSPA5, IL10RA, IL12A, IL1RN, **ITGA5**, **ITGAX**, KCNAB1, LDLR, LPL, LRP8, **MMP2**, **MMP3**, NCF1, PC, PHLDA1, PRKCB, PTGS2, PTP4A3, SERPINA3, SREBF2, **TGFB1**, TIAM1, **TIMP1**, TNIP1, TREM2, UCP2 | -3.727 | 1.88E-10 |
|  | Estrogen receptor | group | ANXA1, BCL2, CAPG, CD44, CD68, CDH11, CDH19, CDH4, CNKSR1, **COL4A1**, **COL4A2**, **COL4A5**, **COL5A1**, **COL6A1**, **COL6A2**, EGFR, EGR1, ERBB2, **FLNC**, FOS, GSTA5, HBEGF, HSPA1A/HSPA1B, KLF10, KRT18, KRT7, KRT8, **LAMC2**, LDLR, **LOXL2**, MAP1B, **MMP14**, MSN, PCDH7, PLAUR, RGS2, **TGFB1**, **TIMP1**, TLR4, TNC, **VIM** | -3.661 | 1.34E-07 |
| **Activated** | Platelet-derived growth factor subunit BB  (PDGF BB) | complex | ATF3, BCL3, BMP6, BRCA1, CASP4, CCNE1, CD44, CDK1, CDKN1A, **COL18A1**, **COL3A1**, CRYAB, CTH, DUSP5, EGFR, EGR1, EGR2, EGR3, FASN, FHL1, FOS, FOSB, FOSL1, FZD1, GADD45A, GDF15, GEM, GLUL, GSS, H19, HBEGF, HLA-E, Hmga2, IER2, IGFBP4, **ITGA5**, JUNB, KLF10, LDLR, **LGALS3**, **LMNA**, **MMP12**, **MMP14**, **MMP2**, **MMP3**, Mt1, Mt2, PHLDA1, PLAT, PLK2, **POSTN**, PRRX2, PTGS2, Pvr, RGS1, RGS2, RXRG, S1PR2, Scd2, SERPINA3, SLC2A3, SLC7A1, SPHK1, TEAD4, **TGFB1**, THBS1, **TIMP1**, TNC, TNFAIP3, TNFRSF12A, VCAN | 6.167 | 2.46E-17 |
|  | Colony stimulating factor 2 (CSF2) | cytokine | ADA, **ADAM8**, ADGRE5, **ANXA1**, ATXN1, AURKA, BBC3, BCL2, BCL3, BID, BIRC3, BIRC5, BUB1, BUB1B, CASP3, CCNA2, CCNB1, CCR5, CD14, CD180, CD276, CD63, CD80, CD83, CD86, CDCA2, CDCA8, CDK1, CDKN1A, CENPE, CIITA, CKS1B, CLEC6A, CLEC7A, **COL8A1**, CXCL10, CYBB, E2F8, EGR1, EGR2, EGR3, F2R, F2RL1, FIGNL1, FOS, FOSL1, FOXM1, GDF15, HBEGF, HGF, IL1RN, **ITGAX**, JAK2, JUNB, KIF11, LCP1, LY9, LY96, MKI67, **MMP14**, **MMP2**, NFKB2, PRC1, PTGER2, PTGS2, PTK2B, QSOX1, REC8, RELB, RRM2, SLAMF7, SLC2A1, SLC2A3, SLC2A4, SOCS1, SOCS2, SREBF2, **TGFB1**, THBS1, TICAM1, TLR2, TLR4, TNFAIP3, TOP2A, UHRF1, UPP1 | 6.258 | 2.35E-15 |
|  | Immunoglobulin E (IGE) | antibody | **ADAM8**, **ANXA1**, ASB2, BAIAP2, BCAT1, BCL2, BCL3, BIRC5, CAPN2, CCL22, CCL5, CCR5, Cd33, CD80, CD86, CDK19, CDKN1A, CLEC7A, **COL18A1**, CTSK, CX3CL1, CXCL16, DUSP2, DUSP4, EGR1, EGR2, EMILIN1, ENO2, ERRFI1, F2R, FYN, GADD45B, GDF15, HAVCR2, HBEGF, HIP1R, HIVEP3, IL7R, **ITGA5**, **ITGAV**, **ITGAX**, JAK2, JUNB, LRRC38, MDFIC, NCF4, NFKB2, NFKBIE, PDGFB, PILRA, Plpp1, PTGS2, PTPN6, PXMP2, RAI14, RASGRP1, RELB, **RHOD**, RNF180, RUNX1, SERPINB1, SERPINB6, Serpinb6b, SLC11A1, SLC37A2, SOCS1, SPHK1, SRGAP3, STAT5A, **TGFB1**, TLR2, TNC, TNFRSF10A, TNFRSF11B, TNFRSF12A, TNFRSF13B, Tnfrsf22/Tnfrsf23, TRAF1, **TUBB6**, UGCG, **VIM** | 6.407 | 1.9E-23 |
|  | Transforming Growth Factor Beta 1 (TGFB1) | growth factor | ABI2, ACTC1, **ADAM19**, **ADAMTS12**, **ADAMTS3**, **ADAMTS4**, ADI1, ADK, ALDH18A1, ALOX12, AMD1, ANGPTL4, **ANKRD1**, ANPEP, **ANXA2**,  **ANXA8/ANXA8L1**, ASS1, ATXN1, B3GALT2, BBC3, BCL2, BCL3, BDH1, BGN, BIRC5, BMP6, BUB1, BUB1B, C1QA, C3AR1, C5, Calm1, CASP3, CASP4, CBR3, CCL5, CCNA2, CCNB1, CCNB2, CCNE1, CCR5, CCRL2, CD14, CD4, CD44, CD68, CD72, CD80, CD83, CD86, CDH11, CDH19, CDH4, CDK1, CDK5R1, CDKN1A, CDKN2A, CDT1, CELSR2, CENPE, CENPF, CIITA, CKS1B, CLIC4, CLU, **COL18A1**, **COL3A1**, **COL4A1**, **COL4A2**, **COL5A1**, **COL6A1**, **COL6A2**, **COL6A3**, **COL8A1**, COTL1, CPXM1, CSPG4, **CTSB**, **CTSK**, **CTSS**, CTTN, CX3CL1, CX3CR1, CXCL10, CXCR6, CYBB, DAPK1, DBP, DKK3, DUSP4, EDNRA, EEF1A1, EGLN1, EGR1, EGR2, EGR3, EIF4EBP1, ELF3, EMILIN1, ENO2, ESPL1, F2R, F2RL1, FAM110B, FASN, FETUB, FGFBP1, FHL1, FNDC5, FOS, FOSB, FYN, FZD1, GADD45A, GADD45B, GCNT1, GDF15, GEM, GPRC5B, GSDME, GSTA5, HBEGF, HDAC9, HEXA, HGF, HK1, HMGA1, HNMT, HSF2BP, HSPA1A/HSPA1B, HSPA5, HSPB1, IER2, IFI30, IFIT3, IGFBP3, IGFBP4, IGHM, IL10RA, IL12A, IL1RN, IL2RB, IRAK2, **ITGA5**, **ITGAV**, **ITGAX**, **ITGB2**, ITIH5, JUNB, KCNG1, KDELR3, KLF10, KLF15, KLK3, KRT18, KRT7, KRT8, **LAMC2**, LDLR, **LGALS3**, LIMS1, LITAF, LOC102724788/PRODH, **LOXL1**, **LOXL2**, LPL, MAOA, MBOAT2, ME2, MFAP2, MGAT5, MKI67, **MMP12**, **MMP14**, **MMP2**, **MMP3**, MSMO1, MSN, MSTN, **MYL3**, MYOG, NAB2, NCAM1, NCF1, NDRG4, NEGR1, NPAS2, PAPPA, PARP3, PDGFB, PDPN, PILRA, PLAT, PLAUR, PLK2, PLXNC1, PMM1, **POSTN**, PPT1, PRC1, PROM1, PSAT1, PTGER2, PTGS2, PTK2B, PTP4A3, PTPN6, RAB31, RASGRP1, **RHOC**, **RHOD**, RIN1, RND1, RNH1, RRAD, RUNX1, RUNX2, RUNX3, S100A10, S100A4, S1PR2, SAR1B, SBNO2, SELENBP1, SELPLG, SERPINA3, SERPINB1, SHMT1, SLC16A9, SLC1A2, SLC2A1, SLC2A3, SLC7A1, SOCS1, SOX4, SOX9, SPHK1, SSTR2, STAT5A, STAT5B, TAB2, TFAP4, **TGFB1**, TGIF1, THBS1, **TIMP1**, TLR2, TLR4, TMIGD1, TNC, TNFAIP3, TNFRSF10A, TNFRSF11B, TNFRSF12A, **TNNT2**, TOP2A, TP73, TRAF1, TRIM9, **TUBA1A**, **TUBB2A**, UCK2, ULK1, USH1C, VAT1, VCAN, **VIM**, WISP1, WNT11, WNT4, ZFP36L2, ZFPM2, ZNF365 | 6.738 | 6.92E-35 |
|  | Tumor necrosis factor (TNF) | cytokine | A4GALT, ABR, ACADM, **ADAM8**, **ADAMTS4**, **ADAMTS7**, **ADAMTS8**, AGT, AKR1B10, AMPD3, ANGPTL4, ANPEP, **ANXA1**, ARC, ARHGAP22, ARL6IP5, ASS1, ATF3, B4GALNT1, BBC3, BCKDHA, BCKDHB, BCL2, BCL3, BGN, BID, BIRC3, BIRC5, BMPER, BPGM, BTG2, BUB1B, C3AR1, C5, CA2, CASP3, CASP4, CBR3, CCL22, CCL28, CCL5, CCNE1, CCR5, CD14, CD4, CD44, CD80, CD82, CD83, CD86, CDH11, CDK5R1, CDKN1A, CDKN2A, CERS6, CHRND, CHSY3, CIB2, CIITA, CLEC11A, CLIC4, CLU, CNR2, **COL15A1**, **COL27A1**, **COL3A1**, COLQ, COTL1, CRYAB, **CTSB**, **CTSK**, **CTSS**, **CTSZ**, CTTN, CX3CL1, CX3CR1, CXCL10, CXCL16, CYBB, CYP27A1, CYTIP, DBT, DLL4, **DMD**, DUSP2, DUSP4, DUSP5, EFHD2, EGFR, EGLN1, EGR1, EGR2, EGR3, ELF3, EMP1, EMP2, ERBB2, EXOC3L4, F2RL1, FADD, FASN, FOS, FOSB, FOSL1, FOXO4, FST, FUT4, FYN, GABRA1, GABRG2, GADD45A, GADD45B, GBP6, GDF15, GEM, GPD1, GPD2, GPRC5B, GPX1, GRN, H19, HBEGF, HDAC9, HEXA, HGF, HID1, HLA-A, HLA-E, HPCA, HPGDS, HSPA1A/HSPA1B, HSPA5, IER2, IFIT3, IGFBP3, IGFBP4, IL10RA, IL12A, IL1RN, IL21R, IL7R, IRAK2, IRS2, **ITGA5**, **ITGAV**, **ITGAX**, **ITGB2**, JUNB, KIF20A, KLF10, KLF5, KLK3, KRT18, KRT8, **LAMC2**, LDLR, **LGALS3**, LITAF, LPL, LY96, MAP2K6, MAP3K14, MEOX1, MFHAS1, **MMP12**, **MMP14**, **MMP16**, **MMP2**, **MMP3**, MSLN, MSTN, Mt1, Mt2, MVP, MYOG, NCAM1, NCF1, NFKB2, NFKBIE, NOD2, NQO1, OASL, P2RX5, PAPPA, PC, PCDH7, PDGFB, PDIA4, PDK3, PDPN, PER2, PHLDA1, PLA2G16, PLA2G5, PLAT, PLAUR, PLD3, PLK2, PLXNB2, **POSTN**, PTGS2, PXMP2, PYCARD, RASSF7, RELB, RGS1, RGS2, RND1, RRAD, RRM2, RUNX2, SCUBE2, SELPLG, SERPINA3, SERPINB1, SERPINB8, SGPL1, SLC11A1, SLC15A3, SLC16A2, SLC1A2, SLC22A4, SLC2A1, SLC2A4, SLC40A1, SLC7A1, SLC7A2, SNN, SOAT1, SOCS1, SOCS2, SOX4, SOX9, SPHK1, SQLE, STAT5A, TALDO1, TBXAS1, TFPI2, TGFB1, TGIF1, Tgtp1/Tgtp2, THBS1, TICAM1, **TIMP1**, TLR2, TLR4, TMEM176B, TNC, TNFAIP3, TNFRSF10A, TNFRSF11B, TNIP1, TP63, TRAF1, TREM2, TST, TUB, TYK2, UACA, UCP2, UGCG, USP2, **VIM**, WISP1, ZNF365, ZNF750 | 7.785 | 6.19E-32 |

Supplementary Table 3. Ingenuity Pathway Analysis – RNAsequencing *mdx* vs *mdx* ^TG^

*Target genes in red are ECM, ECM-associated, cytoskeletal, or cytoskeletal-associated genes.

| **Predicted Activation State** | **Upstream Regulator** | **Molecule Type** | **Target Genes** | **Activation z-score** | **p-value of overlap** |
| --- | --- | --- | --- | --- | --- |
| **Inhibited** | Prostaglandin E Receptor 4 (PTGER4) | G-protein coupled receptor | Ccl7, CXCL10, CXCR4, EGR1, GDNF, GLIS3, HIVEP3, IGF2BP2, PDGFB, **SPP1** | -2.345 | 0.00765 |
|  | Dachshund Family Transcription Factor 1 (DACH1) | transcription regulator | CDKN1A, EGR1, IER2, IGFBP3, TNFAIP3 | -2.236 | 0.00329 |
|  | Scaffold Attachment Factor B (SAFB) | Nuclear matrix | BBC3, CNTNAP2, CX3CL1, CXCL10, HEXA | -2.219 | 0.0146 |
|  | Prostaglandin E Receptor 2 (PTGER2) | G-protein coupled receptor | AURKA, CENPE, CLEC4D, CXCR4, EGR1, ESD, **ITGAL**, MELK, **SPP1** | -2.138 | 0.00207 |
|  | SP110 Nuclear Body Protein (SP110) | transcription regulator | CLU, CXCL10, EGR3, MAOA, PANX1, PERP, SOX4, SQSTM1 | -2.121 | 0.0212 |
| **Activated** | Protein Kinase A (Pka) | complex | ARC, CDKN1A, CXCL10, CYP51A1, DUSP4, EGR1, JUNB, SLC1A2, SOX9 | 2.975 | 0.0145 |
|  | Transforming Growth Factor Beta 1 (TGFB1) | growth factor | ABCG1, ABI2, ALOX12, AMD1, ASS1, BBC3, C5, CBR3, Ccl7, CD300A, CD68, CDKN1A, CENPE, CLU, CSPG4, CTTN, CX3CL1, CXADR, CXCL10, CXCR4, CXCR6, DEPTOR, DUSP4, EDNRA, EGF, EGR1, EGR2, EGR3, EIF4EBP1, ELF3, ENO2, F2RL1, FABP5, FETUB, FGFBP1, FNDC5, GAL, GDF15, GDNF, GSDME, GSTA5, HEXA, HK1, HMGA1, HSF2BP, HSPA1A/HSPA1BID4, IER2, IER3, IGF2, IGFBP3, IGFBP5, IGHM, IL12A, IRAK2, **ITGAL**, JUNB, KRT7, **LGALS3**, LOC102724788/PRODH, MAOA, Masp1, ME2, MEFV, **MMP12**, MSTN, NAB2, NCAM1, NDRG4, NEGR1, PDGFB, PLA1A, PLK2, PMM1, **RHOC**, RIN1, RNH1, S100A10, S1PR2, SLC16A9, SLC1A2, SLC2A1, SLC2A3, SOX4, SOX9, SPP1, SSTR2, TGIF1, TLR2, TMIGD1, TNFAIP3, TNFRSF12A, TP73, TRIM9, USH1C, VAT1, VCAM1, WNT4, WNT5A | 3.085 | 4.71E-12 |
|  | KRAS proto-oncogene (KRAS) | GTPase | CDKN1A, CLU, CRYAB, CX3CL1, CXADR, CXCL10, DUSP4, DUSP5, EGR1, EGR2, F2RL1, FKBP11, FOSL1, GLUL, HEXA, HMGA1, IER3, IGF2, IGF2BP2, JUNB, KCNN4, **LGALS3**, **LMNB1**, LRP8, LYZ, MSLN, NCAM1, NETO2, NFKB2, NGEF, NQO1, PLA1A, **RHOC**, RNH1, SOX9, **SPP1**, SQSTM1, STOM, TNFRSF12A, UPP1, ZFAND2A | 3.101 | 5.18E-09 |
|  | palmitic acid | chemical - endogenous mammalian | BBC3, CD68, CDKN1A, CPE, CXCL10, CYP4F12, ELOVL6, F2RL1, GDF15, IRAK2, SQSTM1, TLR2, TRIM63, UCHL1, VCAM1 | 3.132 | 0.000965 |
|  | Uncoupling Protein 1 (UCP1) | transporter | AMD1, Cd24a, CD68, CPNE2, CXCL10, EIF4EBP1, ENHO, GDF15, MSS51, ODF3L2, PGD, REEP6, SMOX, SMYD2, STAB2, TPPP, USH1C | 3.85 | 2.11E-06 |
